# Protective immune trajectories in early viral containment of non-pneumonic SARS-CoV-2 infection

**DOI:** 10.1101/2021.02.03.429351

**Authors:** Kami Pekayvaz, Alexander Leunig, Rainer Kaiser, Sophia Brambs, Markus Joppich, Aleksandar Janjic, Oliver Popp, Vivien Polewka, Lucas E. Wange, Christoph Gold, Marieluise Kirchner, Maximilian Muenchhoff, Johannes C Hellmuth, Clemens Scherer, Tabea Eser, Flora Deák, Niklas Kuhl, Andreas Linder, Kathrin Saar, Lukas Tomas, Christian Schulz, Wolfgang Enard, Inge Kroidl, Christof Geldmacher, Michael von Bergwelt-Baildon, Oliver T. Keppler, Ralf Zimmer, Philipp Mertins, Norbert Hubner, Michael Hölscher, Steffen Massberg, Konstantin Stark, Leo Nicolai

**Author notes:** Contributed equally (Kami Pekayvaz, Alexander Leunig). Contributed equally (Konstantin Stark, Leo Nicolai).

## Abstract

The immune system of most SARS-CoV-2 infected individuals limits viral spread to the upper airways without pulmonary involvement. This prevents the development of pneumonic COVID-19. However, the protective immunological responses causative of successful viral containment in the upper airways remain unclear. Here, we combine longitudinal single-cell RNA sequencing, proteomic profiling, multidimensional flow cytometry, RNA-Seq of FACS-sorted leukocyte subsets and multiplex plasma interferon profiling to uncover temporally resolved protective immune signatures in non-pneumonic and ambulatory SARS-CoV-2 infected patients.

We compare host responses in a high-risk patient population infected with SARS-CoV-2 but without pulmonary involvement to patients with COVID-19 pneumonia. Our data reveal a distinct immunological signature of successful viral containment, characterized by an early prominent interferon stimulated gene (ISG) upregulation across immune cell subsets. In addition, reduced cytotoxic potential of Natural Killer (NK) and T cells, as well as a monocyte phenotype with immune-modulatory potential are hallmarks of protective immunity. Temporal resolution across disease trajectories highlights ISG upregulation as particularly prominent early in the disease and confirms increased expression also in comparison to healthy controls.

We validate this distinct temporal ISG signature by in-depth RNA-seq of FACS-sorted leukocyte subsets in a large prospective ambulatory SARS-CoV-2 infected cohort confirming early and robust ISG upregulation particularly in monocytes and T cells. In conclusion, our data demonstrate a protective ISG phenotype in patients with successful containment of SARS-CoV-2 infection without progression to COVID-19. This early protective interferon response might be exploited as a therapeutic approach and for disease course prediction.

## Introduction

Due to its high contagiousness and an approximate case fatality rate of 1.0-2.3% the COVID-19 pandemic has confronted the world with a major health care and economic challenge^1-3^. Approximately 10% of cases have a severe disease course^4^. Extensive data now underlines immunopathology, the concept of self-inflicted damage to tissues by the immune system, as an important factor contributing to disease progression in COVID-19^4-8^.

Various detrimental pathways have been identified to drive immunopathology in COVID-19: A cytokine storm with high levels of IL-6 with some similarity to CAR-T cell hyperinflammation was postulated^9, 10^. Recent data highlighted a failure in host interferon I and III responses associated with progression to severe COVID-19^11-13^. In contrast to influenza, severe COVID-19 has also been associated with impaired Interferon-γ expression levels in T cells^4, 7, 14^. Seminal work revealed auto-antibodies against interferon pathway related proteins and inherent dysfunctional mutations in a significant proportion of severe COVID-19 cases^15, 16^. T cells in severe cases show upregulation of exhaustion and apoptosis markers^5^. Apart from adaptive immunity, the concept of a dysregulated innate immune cell axis evolved consistently across studies, with an expansion of HLA-DR^low^ monocytes and a surge in (premature) neutrophils, which in turn cause vascular inflammation and immunothrombosis in the lung and remote organs^17-20^.

Interestingly, most infected patients, even those with known risk factors like hypertension or obesity, efficiently clear SARS-CoV-2 without developing pneumonia ^21, 22^. This highlights the ability of the host immune system to mount an effective response and contain the virus in the upper airways without causing pulmonary damage in the majority of cases^23^.

Yet, the pathways that provide protective immunity in SARS-CoV-2 infection are less well understood. Outpatients and patients with oligo- and asymptomatic SARS-CoV-2 infection, who represent the exemplary immune response, have been underrepresented in mechanistic studies.

To better understand how protective immune response and immunopathology differ in SARS-CoV-2 infection, we launched a two-cohort multi-omics study characterizing the immune response in ambulatory and non-pneumonic infected individuals. With this experimental approach, we first used a well characterized, exploratory cohort of high-risk patients for hypothesis generation and subsequently confirmed our findings in a large cohort of outpatient SARS-CoV-2 infected individuals. A multi-omics approach allowed us to identify protective cell responses in these individuals by contrasting them with COVID-19 pneumonia cases and healthy controls in a longitudinal manner.

We identify a distinct immunological signature of successful viral containment, featuring a prominent, early interferon stimulated gene (ISG) upregulation across immune cell subsets which is independent of plasma interferon levels. In addition, reduced cytotoxic potential of Natural Killer (NK) and T cells, as well as a monocyte phenotype with immune-modulatory properties were hallmarks of protective immunity.

## Results

### Longitudinal clinical and cellular characteristics of the successful and failing immune response to SARS-CoV-2

We compared high risk patients with an immune response that successfully prevents pulmonary involvement in SARS-CoV-2 infection to the response of patients developing COVID-19 pneumonia. We performed single cell RNA sequencing (sc-RNA seq), in-depth RNA sequencing of sorted immune cell populations, 50-dimensional flow cytometry, plasma shotgun proteomics together with cytokine profiling longitudinally throughout the disease course. We used a longitudinally sampled exploratory high-risk cohort for hypothesis generation (n=14 patients) and a longitudinally sampled outpatient confirmation cohort to validate our results (n=58 patients) (see Methods). For the exploratory cohort, the three sampling time points were after first positive PCR test (TP1), on average five days after initial sampling (TP2) and after negative PCR/seroconversion or symptomatic improvement (TP3) (Figure 1a-b Suppl Fig 1a-b and Methods).

**Fig 1.**
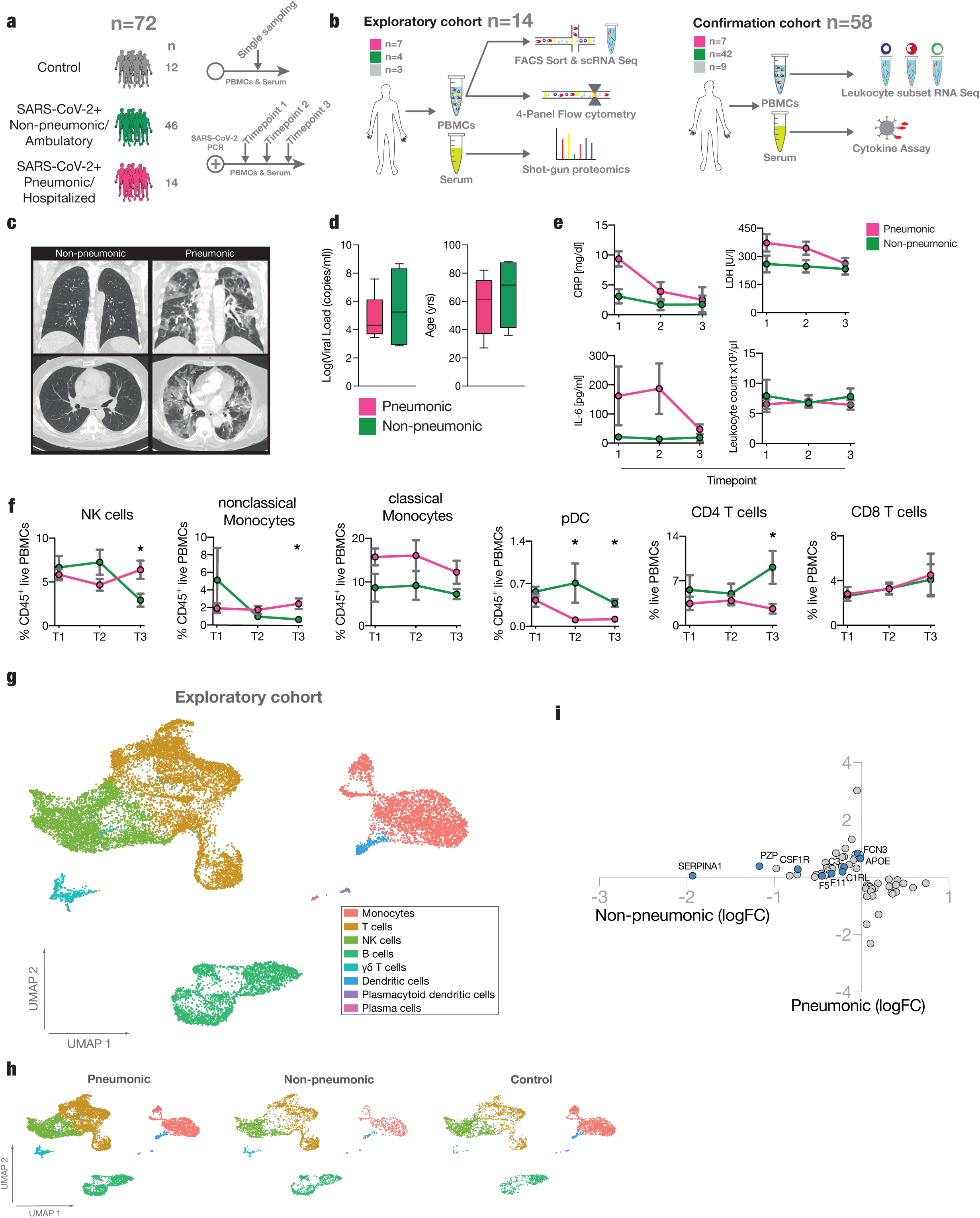
Study overview and longitudinal clinical and cellular dynamics in non-pneumonic and pneumonic SARS-CoV-2 infection. **a-b** Experimental setup and processing pipeline. **a:** Plasma and PBMCs of n=12 Controls, n=14 pneumonic COVID-19, n=46 non-pneumonic/ambulatory infected patients were sampled, in case of pneumonic and non-pneumonic SARS-CoV-2 positive patients at least on three consecutive time points. **b:** Exploratory cohort: longitudinal samples from 11 pneumonic and non-pneumonic infected patients and one timepoint from three control patients were used for shotgun plasma proteomics, 50-dimensional flow cytometry or single cell RNA sequencing. Confirmation cohort: Longitudinal samples from a total of 42 ambulatory COVID-19 patients and one sample from n=9 controls and n=7 hospitalized COVID-19 patients were used for cytokine measurements, or cell sorting subsequent bulk RNA sequencing. **c:** Representative axial and coronal computed-tomographic scans of hospitalized pneumonic and non-pneumonic infected patients. **d:** Baseline characteristics of the exploratory group: log10 of viral load as measured in copies/ml of upper respiratory tract swap samples. n=6 pneumonic COVID-19, n=4 non-pneumonic infected patients. Age in years of patients. n=7 pneumonic COVID-19, n=4 non-pneumonic for both. **e:** Qualitative longitudinal clinical laboratory values of CRP (mg/dl), LDH(U/l), IL-6 (pg/ml) and total leukocyte count (1000/µl) at time points 1-3. n=7 pneumonic COVID-19, n≥3 non-pneumonic COVID-19 patients per time point. **f:** Percentage of immune cells as live CD45^+^ cells/PBMCs measured by flow cytometry per sampling time point. Two-sided t-test, n≥3 per time point and group. **g:** UMAP representation of the pooled exploratory cohort showing the assigned cell populations. **h:** UMAP representation of the sequenced samples showing the assigned cell populations per group for all time points. **i:** Plot depicting significantly (p<0.05) differentially expressed proteins in plasma samples pooled across all three time points. Log fold changes are computed relative to the control proteins’ expression. All error bars are mean ± s.e.m. unless otherwise noted. *p<0.05.

In our exploratory cohort, radiological assessment of non-pneumonic, SARS-CoV-2 infected patients revealed no signs of lung injury on high-resolution computed tomography compared to patients experiencing pulmonary symptoms (Figure 1c). Upper respiratory tract viral loads derived from nasopharyngeal swabs between pneumonic and non-pneumonic patients in the exploratory group were comparable (Figure 1d). Patients in the exploratory cohort had on average 2.9 risk factors for developing a severe disease course, with all patients having at least 1 risk factor for a severe disease course, and a trend towards more risk factors in the non-pneumonic cohort (Suppl. Table 1, see methods).

Absolute leukocyte counts did not differ between groups, while other inflammatory markers such as CRP, IL-6 and Lactate Dehydrogenase (LDH) were elevated in the pneumonic group as expected (Figure 1e). Further differentiation of leukocyte subsets revealed lower lymphocyte counts in the pneumonic group, but similar neutrophil and monocyte counts as reported previously (Suppl. Fig. 1c)^8, 12, 24, 25^.

To allow for an in-depth analysis of leukocyte subsets, we performed phenotyping of surface marker expression by flow cytometry (Suppl. Fig. 1d). We found relative NK-cell counts in non-pneumonic infected patients dropped significantly in comparison to pneumonic patients at timepoint 3 (Figure 1f). Similarly, changes in the mononuclear phagocyte (MNP) compartment included no major differences in monocyte subsets and significantly higher relative pDC counts in non-pneumonic patients compared to patients with pulmonary injury (Figure 1f).

We further observed distinct shifts in T cell populations, mainly presenting with a faster recovery of relative CD4^+^ T cell counts in non-pneumonic patients, but without numerical changes in CD8^+^ T cells (Figure 1f).

Unsupervised UMAP-clustering of single cell RNA-seq data of PBMCs identified 17 distinct immune cell populations (Suppl. Fig. 1e). The subclusters were integrated into the following immune cell populations: T cells, B cells, monocytes, NK cells, myeloid dendritic cells, plasmacytoid dendritic cells, plasma cells and gamma delta T cells (Figure 1g-h). T-cells were further differentiated into CD4^+^ (Population 5), CD8^+^ (Populations 1, 4, 8) and double negative T cells (Population 13). The top cluster defining genes are depicted in (Suppl. Fig. 1f). In line with flow cytometric data, UMAP revealed no striking differences in immune cell composition across disease states and time points (Figure 1h, Suppl. Fig. 2a).

Plasma shotgun proteomics identified 1102 proteins. Internal quality control revealed a strong positive correlation of CRP determined by our proteomic approach with clinical CRP (r = 0.9136) and fibrinogen measurements (r = 0.7967, Suppl. Fig. 2b). Principal component analysis slightly separated control patients from SARS-CoV-2 infected patients, without strong differences between disease severities (Suppl. Fig. 2c). Comparison with another plasma proteome study of COVID patients^26^ showed correlations of the plasma proteome up to 38% (Suppl. Fig. 2d). 171 plasma proteins were significantly upregulated in either pneumonic or non-pneumonic SARS-CoV-2 infection and downregulated in the other in comparison to healthy controls (Suppl. Fig. 2e). Especially acute phase plasma proteins (APO-E, PZP, CSFR1, SERPINA1), coagulation factors (FCN2, F5, F11) and components of the complement system (C3, C1RL) were upregulated in pneumonic COVID-19 patients (Figure 1i, Suppl. Fig. 2f-g).

Taken together, comparison of non-pneumonic and pneumonic patients revealed only mild global changes in circulating immune cells and plasma proteins, consistent with previous reports^18^. To better understand how the cellular response to SARS-CoV-2 infection might shape disease course, we next analyzed our transcriptomic data in detail.

### Enhanced interferon response across immune cell populations defines protective immunity in non-pneumonic SARS-CoV-2 infection

We first aimed to identify protective immune cell signatures associated with non-pneumonic disease course. We compared differentially expressed genes (DEGs) across sampling time points between the two disease states. Assessment of T cell, B cell, NK cell, and monocyte populations revealed consistent upregulation of interferon stimulated genes (ISGs) in the non-pneumonic cohort. For example, transcripts encoding Interferon Induced Protein 44 Like (*IFI44L*) and Interferon-induced Transmembrane Protein 1 (*IFITM1*), both of which are involved in viral containment^27, 28^, showed significantly higher levels in both CD4^+^ T cells and NK cells of non-pneumonic SARS-CoV-2 infected patients (Figure 2a-c). Additionally, interferon signaling in B-cells seemed to be more prominent in patients without pulmonary injury, indicated by increased expression of *MX1*^29^, *XAF1*^30^ and *IFI44L* compared to patients with pneumonic COVID-19 (Figure 2d-e). *IFI44L*^27^, *IFI44*^27^, *IFI6*^31^, *LY6E*^31^, *ISG15*^32^ were significantly higher expressed in monocytes from non-pneumonic cases compared to patients with pulmonary injury (Figure 2f-g). When transforming the transcriptomic data from single cell resolution to bulk-data across all subpopulations, *IFI44L*^27^ and *IFITM1*^23^ were significantly upregulated across all cells in non-pneumonic patients compared to pneumonic COVID-19 patients across all timepoints (Figure 2i).

**Fig 2.**
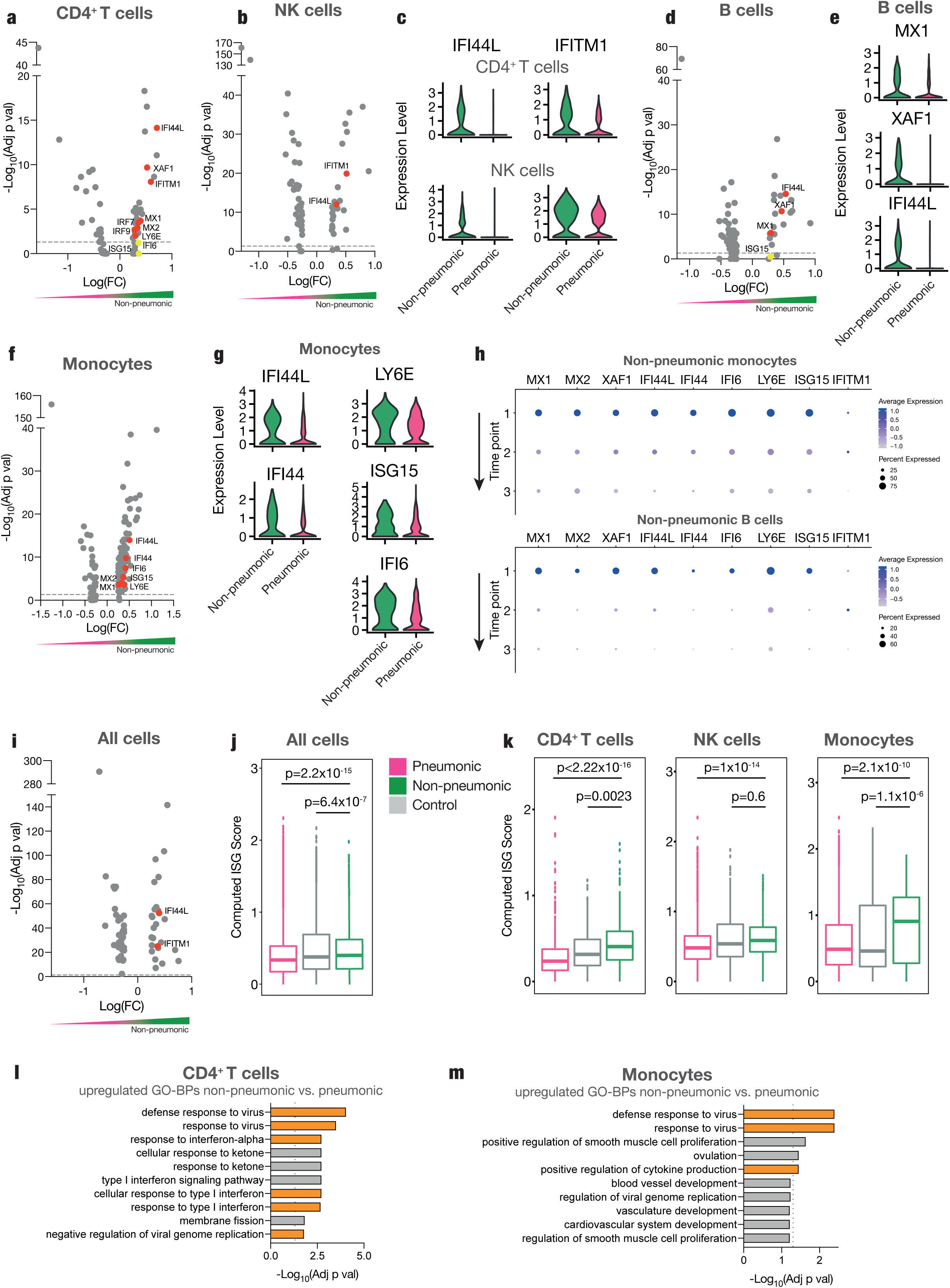
Enhanced interferon response across immune cell populations defines protective immunity in SARS-CoV-2 infection. **a-b:** Volcano plots of differentially regulated genes in CD4^+^ T cells and NK cells of pneumonic compared to non-pneumonic samples. Genes enriched in pneumonic samples have negative log(FC), genes enriched in non-pneumonic samples have positive log(FC). The color scale underneath emphasizes this (red pneumonic, green non-pneumonic) **c:** Violin plots of expression of IFI44L and IFITM1 in CD4^+^ T cells and NK cells for pneumonic and non-pneumonic samples. **d:** Volcano plot of differentially regulated genes in B cells of pneumonic compared to non-pneumonic samples. **e:** Violin plots of expression of MX1, XAF1 and IFI44L in B cells for pneumonic and non-pneumonic samples. **f:** Volcano plot of differentially regulated genes in monocytes of pneumonic compared to non-pneumonic samples. **g:** Violin plots of expression of IFI44L, IFI44, LY6E, ISG15 and IFI6 in monocytes for pneumonic and non-pneumonic samples. **h:** Dot-plot of the scaled average expression and Percent expressing cells of selected interferon stimulated genes in monocytes and B cells by sampling time point in non-pneumonic patients. **i:** Volcano plot of differentially regulated genes in all sequenced cells of pneumonic compared to non-pneumonic samples. **a-b, d, f, i:** Red annotations are significantly upregulated (adj p val<0.05), yellow ones are non-significantly differentially expressed. Positive fold change signifies higher expression in the non-pneumonic group. Line denotes adj p val<0.05. **j:** Box plots of computed ISG-score (see methods) of all sequenced cells. **k:** Box plots of ISG-scores of CD4^+^ T cells, NK cells and monocytes. P values are shown above, non-pneumonic with pneumonic and control with pneumonic are compared. **a-g, i-m**: Longitudinal samples are pooled if not otherwise indicated. **l-m:** Top 10 Gene Ontology - Biological Processes (GO-BPs) terms from upregulated genes of non-pneumonic vs pneumonic samples. Pathways of interest are marked, line shows adj p val<0.05.

When resolving these data temporally, we noticed interferon signaling related genes across different immune cell subsets to be most prominently and most abundantly expressed at the earliest time point during the disease course in non-pneumonic patients (Figure 2h, Suppl. Fig. 3a). This underlines that ISGs might play an early and deterministic role in shaping COVID-19 disease course. *In vitro* studies on SARS-CoV-2 have revealed a suppressed or delayed interferon response by infected cells, mediated by multiple viral proteins inhibiting the physiological antiviral response^33, 34^. We therefore asked if the observed differences simply indicated a down-regulation of the interferon response in severe COVID-19. However, when comparing non-pneumonic SARS-CoV-2 infected patients with uninfected control patients, we found persistent differential expression, pointing towards an active enhancement of ISG signaling in peripheral immune cells as a key immunological feature in non-pneumonic disease (Suppl. Fig.3b, d-f).

To further validate these findings, we computed an ISG-score for each cell population^12^. Indeed, the summarized ISG score across all cell populations revealed that non-pneumonic cases displayed a substantial rise compared to pneumonic patients but also compared to non-COVID-19 controls (Figure 2j). The strong increase in ISGs between pneumonic and non-pneumonic COVID-19 patients was reproducible across CD4^+^ T cells, NK cells and was particularly prominent in monocytes (Figure 2k). Finally, gene ontology (GO) term analysis of biological processes revealed an enrichment of “defense response to virus”, “response to virus”, “response to interferon-alpha” “response to type I interferon”, “negative regulation of viral genome replication” and several other interferon-related pathways in CD4^+^ T cells and monocytes (Figure 2l-m and Suppl. Fig. 3c).

Together, these data point towards a protective global induction of ISGs in peripheral immune cells of patients able to successfully contain SARS-CoV-2 infection.

### Decreased cytotoxic potential of lymphocytes in non-pulmonary SARS-CoV-2 infection

To explore additional protective immune cell trajectories, we analyzed the transcriptome of lymphocytes in greater detail. Cytotoxic lymphocytes are key to both the adaptive and innate immune response to viral infections, with cytotoxic T cells and NK cells representing adaptive and innate antiviral branches respectively^35^. Prior work has highlighted upregulation of cytotoxic pathways in severe COVID-19, which could mediate harmful tissue damage^36, 37^.

However, NK cells of infected patients without lung involvement exhibited a decrease in markers of activation and cytolysis, such as granzyme B (*GZMB*)^27^’ granzyme H (*GZMH*)^38^ as well as galectin-1 and galectin-3 (*LGALS1, LGALS3*)^39^ (Figure 3a-b and Suppl. Fig. 4a), compared to pneumonic controls. NK cell activation markers and neutrophil chemoattractants *S100A9* and *S100A8*^*40*^ were less abundantly expressed in non-pneumonic infected patients. In line, NK cells from pneumonic patients showed an upregulation of GO terms such as “cytolysis”, “granulocyte chemotaxis” and “granulocyte migration” (Figure 3c, Suppl.Fig. 4b). Flow cytometry-based surface profiling revealed five NK cell subpopulations among pneumonic and non-pneumonic patient groups characterized by differential surface expression (Suppl. Fig. 4c-g). Subpopulation NK2, characterized by increased expression levels of CD18 necessary for cytotoxic activity^41^, and CD9, a tetraspanin implicated in innate immune cell activation and diapedesis^42^, was detected at high levels throughout pneumonic disease (Suppl. Fig. 4d, f).

**Fig 3.**
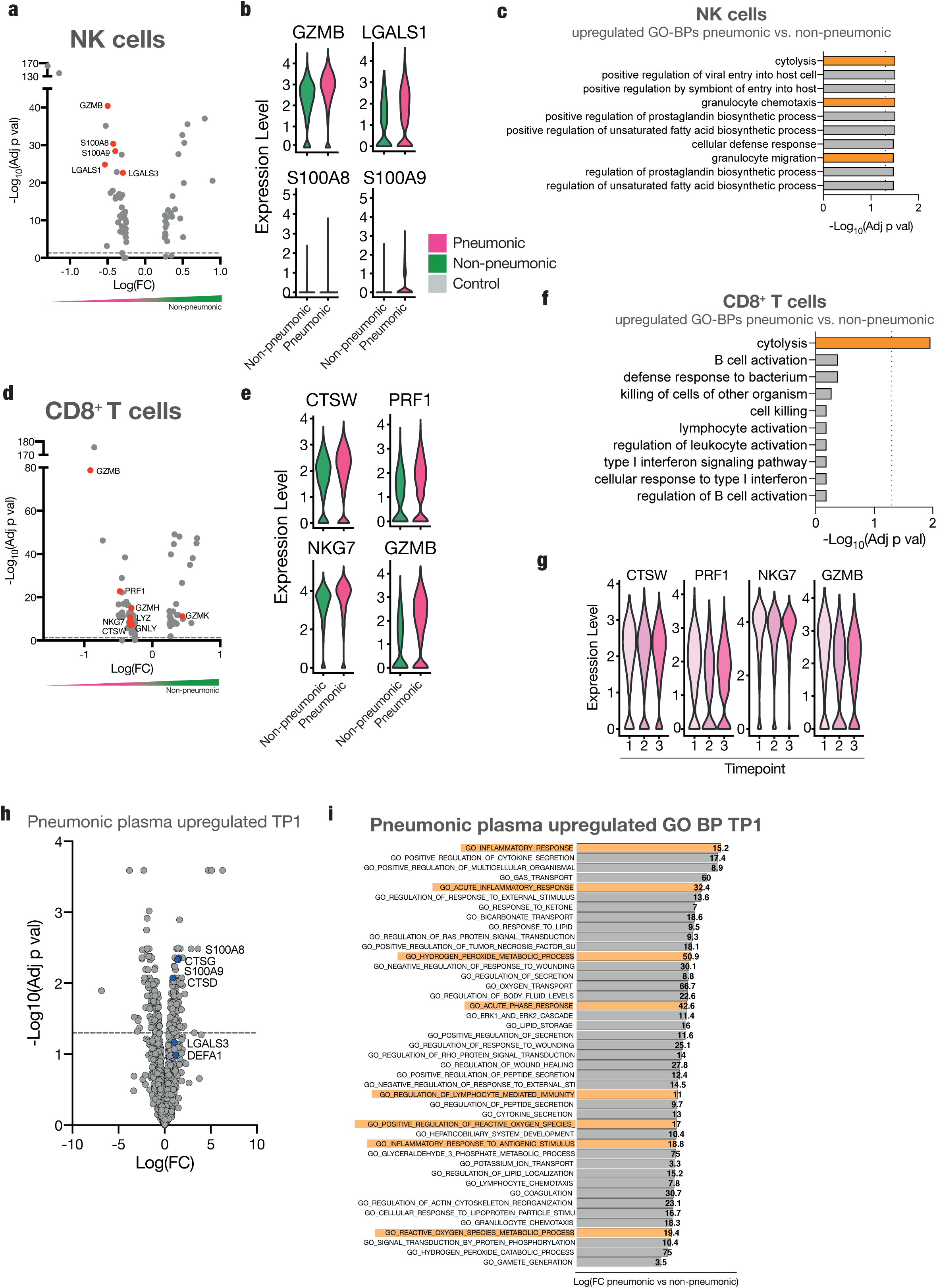
Decreased cytotoxic potential in lymphocytes in non-pneumonic SARS-CoV-2 infection. **a:** Volcano plot of differentially regulated genes in NK cells of pneumonic compared to non-pneumonic samples. **b:** Violin plots of expression of GZMB, S100AB, LGALS1 and S100A9 in NK cells for pneumonic and non-pneumonic samples. **c:** Top 10 GO-BPs from upregulated genes of non-pneumonic vs pneumonic samples in NK cells. Pathways of interest are marked, line shows adj p val<0.05. **d:** Volcano plot of differentially regulated genes in CD8^+^ T cells of pneumonic compared to non-pneumonic samples. **a**,**d:** Red annotations are significantly differentially regulated (adj p val<0.05. Positive fold change signifies higher expression in the non-pneumonic group. Line denotes adj p val<0.05. **e:** Violin plots of expression of CTSW, PRF1, NKG7 and GZMB in CD8^+^ T cells for pneumonic and non-pneumonic samples. **f:** Top 10 upregulated GP-BPs of non-pneumonic vs pneumonic samples in CD8^+^ T cells. Pathways of interest are marked, line shows adj p val<0.05. **g:** Violin plots of expression of CTSW, PRF1, NKG7 and GZMB in CD8^+^ T cells per sampling time point. **h:** Volcano plot of differentially expressed plasma proteins at TP1 of pneumonic samples compared to control samples. Line denotes adj p val<0.05. **i:** Top significantly enriched GO-BPs for pneumonic plasma proteins compared to non-pneumonic plasma proteins. Pathways of interest to lymphocyte cytotoxicity are marked.

Similarly, infected patients without lung injury showed a less cytotoxic CD8^+^ T cell transcriptomic phenotype with decreased expression levels of cytotoxic genes such as *CTSW*^38, 43^, *PRF1*^44^, *NKG7*^40^, *GZMB*^45, 46^, *GZMH*^38^ or *GNLY*^47^ (Figure 3d-e). Accordingly, GO overrepresentation analysis for biological processes revealed a downregulation of the cytolysis pathway in CD8^+^ T cells from patients without pulmonary injury (Figure 3f, Suppl. Fig. 4h). Temporal resolution revealed the respective cytotoxicity genes in CD8^+^ T cells and NK cells to be expressed early in pneumonic COVID-19 patients, however with still a high expression throughout the disease course (Figure 3g).

Plasma proteomic analyses were in line with decreased cytotoxic features of NK and T cells in non-pneumonic infection. *S100A8, S100A9* and *LGALS3* were enriched in pneumonic plasma, especially during the first time point, whereas non-pneumonic patients did not have an enrichment of these markers (Figure 3h, Suppl. Fig. 4j). These markers have previously been shown to be upregulated in severe COVID-19 and are associated with a severe disease outcome^48^. Other indicators of the increased cytotoxicity, cathepsin G and D, and defensin alpha 1, which is not only expressed by neutrophils, but also NK cells, were upregulated at multiple sampling timepoints (Figure 3h, Suppl. Fig. 4j)^49, 50^. Pathway analyses of differentially expressed plasma proteins also revealed upregulated inflammation, acute response and oxygen species pathways, with the most pronounced upregulating at the first sampling point (Figure 3i, Suppl. Fig. 4k). Indeed, cytotoxic T lymphocytes and NK cells have been shown to mediate cytotoxicity by release of reactive oxygen species^51^.

In summary, elevated levels of cytotoxicity in lymphocytes seem to be dispensible for an effective upper-airway containment in non-pneumonic SARS-CoV-2 infection.

### Abundant naïve and immune-modulatory T-cells and antiviral NK-cells in non-pneumonic SARS-CoV-2 infection

In-depth analysis of CD4^+^ T cells revealed higher counts of antigen inexperienced CD45RA^+^ CD4^+^ T cells in non-pneumonic patients, most pronounced at late stages (Figure 4a). In line, antigen experienced CD45RO^+^ CD4^+^ T cells diverged significantly during the disease course in non-pneumonic compared to pneumonic patients (Suppl. Fig. 5a).

**Fig 4.**
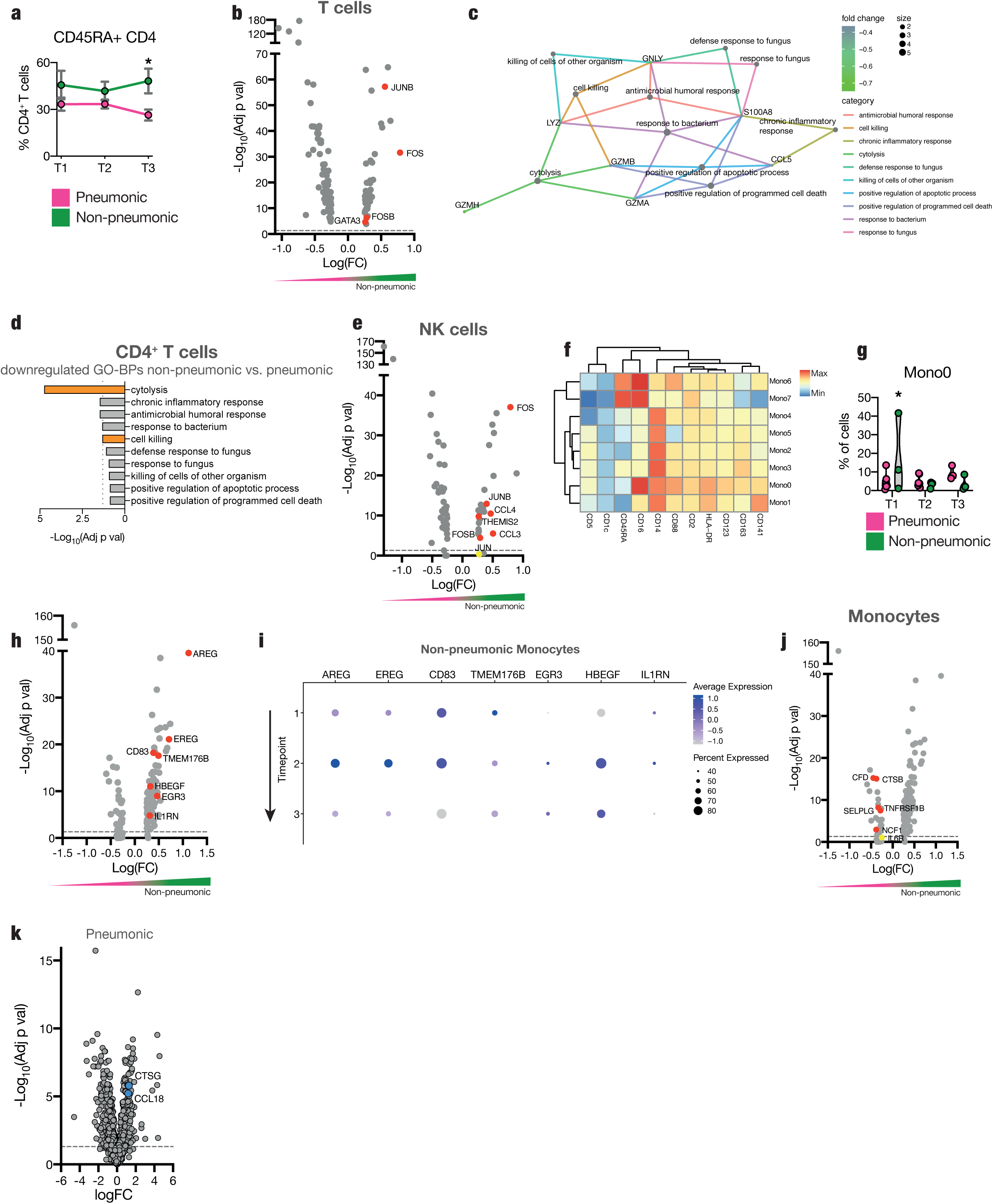
Abundant unexperienced and immune-modulatory T-cells and antiviral NK-cells and anti-inflammatory and anti-thrombotic monocytes in non-pneumonic SARS-CoV-2 infection. **a:** Percentage of CD45RA^+^ CD4^+^ T cells of live PBMCs measured by flow cytometry per sampling time point. Two-sided t-test, n≥3 per time point and group. **b:** Volcano plot of differentially regulated genes in all T cells of pneumonic compared to non-pneumonic samples. **c-d:** GO-BP network analysis and Top10 downregulated GP-BPs of non-pneumonic vs pneumonic samples in CD4^+^ T cells. Pathways of interest are marked, line shows adj p val<0.05. **e:** Volcano plot of differentially regulated genes in NK cells of non-pneumonic compared to pneumonic samples. **f:** Heat map of surface marker expression by FlowSOM group of flow cytometric measurement of monocytes. **g:** Violin plot of Mono0 FlowSOM cluster per time point. Mixed-effects model analysis. Post-hoc Sidak’s multiple comparisons test for individual significant differences between pneumonic and non-pneumonic samples per time point. n≥3 per time point and group. **h:** Volcano plot of differentially upregulated genes in monocytes of non-pneumonic compared to pneumonic samples **i:** Expression of selected genes from **h** in monocytes by sampling time point. **j:** Volcano plot of differentially regulated genes in monocytes of non-pneumonic compared to pneumonic samples. **k:** Volcano plot of differentially expressed plasma proteins of pneumonic samples (all time points pooled) compared to control samples (one-sample moderated t-test). Line denotes adj p val<0.05. **b**,**e**,**h**,**j:** Red annotations are significantly upregulated adj p val<0.05. Positive fold change signifies higher expression in the non-pneumonic group. Line denotes adj p val<0.05. All error bars are mean ± s.e.m. *p<0.05.

tSNE analysis with subsequent FlowSOM subclustering of T cell panel 1 revealed a distinct CD4^+^ CD45RA^+^ CCR10^int^ subpopulation (TC6) to be significantly enriched in non-pneumonic patients, with an increased divergence over the disease course (Suppl. Fig. 5b-f). TC6 was further characterized by particularly low expression of surface activation and exhaustion markers, including CXCR3, PD-1 and CTLA-4^52^ (Suppl. Fig. 5c). Interestingly, CXCR3, which is rapidly induced upon activation of naïve T cells, was expressed only at low levels in TC6^53^ (Suppl. Fig. 5c). Further characterization by T cell panel 2 revealed a CD4^+^ CD45RA^+^ antigen inexperienced T cell population (TC0) specific to non-pneumonic patients, similar to TC6 in T-cell panel 1. CD197 (CCR7) and CD27 levels in this population were intermediate, underlining a rather naïve phenotype^54, 55^ (Suppl. Fig. 5g-k). In line with these distinct changes in surface expression, our scRNA-seq data revealed a more immune-modulatory phenotype of T cells in non-pneumonic infected patients. *JUNB* and *FOS* -both enhanced in T cells from non-pneumonic patients-are subunits of the activating protein-1 (AP-1) ^56, 57^, implicated in T-cell differentiation into effector Th2 ^58^. *GATA3*, a master transcription factor for the differentiation of Th2 cells, was similarly upregulated ^59^. This underlines a potentially immunomodulatory or anti-inflammatory phenotype of T cells in non-pneumonic infection^60^. In line, and similar to CD8^+^ T cells, CD4^+^ T-cells from non-pneumonic SARS-CoV-2 infected patients showed downregulation of “cell killing”, “chronic inflammatory response” and “cytolysis” GO terms (Figure 4b-d).

Corresponding to the expression profiles of T cells, the antiviral transcription factors *JUNB, FOS* and *FOSB* – previously shown to govern IFNγ production in NK cells^61^ – were increasingly expressed in NK cells of non-pneumonic patients (Figure 4e). Subcluster NK3, characterized by high L-selectin (CD62L) expression and reminiscent of an antiviral CD56^dim^ NK cell subset^62^, showed a significant increase among non-pneumonic SARS-CoV-2-infected individuals in flow cytometry (Suppl. Fig. 4e-g). This population is capable of producing significant amounts of IFNγ and is involved in multiple antiviral tasks after restimulation and terminal differentiation^62^. Temporal analysis showed that the increased prevalence of these immunomodulatory/anti-inflammatory pathways was constant across acute disease (TP1 and TP2) (Suppl Fig. 6a), lacking the striking early expression peak identified for ISGs (compare Fig 2h).

### Monocytes with immune-modulatory potential in non-pneumonic COVID-19

As severe COVID-19 immunopathology precedes effective adaptive immune cell function, research has highlighted an innate immune axis, particularly monocytes, in COVID-19 immunopathology^8, 48, 63^.

What are the features defining monocytes in patients able to contain the virus in the upper airways? Flow cytometry revealed a population of non-classical monocytes characterized by surface marker expression of CD16 and CD88, as well as HLA-DR particularly early in disease (Mono0), (Figure 4f-g, Suppl. Fig. 6b-e). To get a better insight into functional changes of monocyteNon-classical monocytes have been implicated as crucial immune modulatory and possibly antigen-presenting cells in inflammation and infection^64^. Along these lines, CD83, which limits cytotoxic T cell effector function, was elevated on transcriptional level in monocytes derived from non-pneumonic patients (Figure 4h)^63, 65^. Moreover, anti-inflammatory genes like *TMEM176B*, known to inhibit the inflammasome and therefore IL1-beta production, as well as Interleukin 1 receptor antagonist (*IL1RN*), showed enhanced transcription in monocytes of non-pneumonic infection (Figure 4h)^66^.

Finally, epidermal growth factor receptor (*EGFR*) ligands, which control tissue repair and regeneration, as well as mucosal immunity, were found to be significantly enhanced in non-pneumonic cases (Figure 4h)^67^. In particular, Ampheregulin (*AREG*), Epiregulin (*EREG*), Early growth response protein 3 (*EGR3*), and Heparin Binding EGF Like Growth Factor (*HBEGF*) showed significant upregulation (Figure 4h). *AREG* has been shown to modulate T reg and Th2 function^68, 69^, while *EREG* is crucial for epithelial integrity and resolution of inflammation^70^. Longitudinal analysis revealed that the identified pathways were most prominently upregulated at TP2, pointing at a pro-resolving and immune-modulatory monocyte phenotype developing later over the disease course (Fig. 4i).

In contrast, in patients developing COVID-19 pneumonia *TNFRSF1B* and *IL6R*, genes encoding IL-6 receptor and TNF receptor, were upregulated in pneumonic COVID-19. In addition, effector proteins like lysosomal cysteine protease Cathepsin B were significantly enhanced (Figure 4j). Monocytes have also been implicated in contributing to immunothrombosis in COVID-19^71^. Indeed, in addition to complement factors upregulated in the plasma (Figure 1i), complement factor D and platelet binding PSGL-1 were upregulated in monocytes of pneumonic patients (Figure 4j). Plasma proteomics showed increased pro-inflammatory CCL18, as well as Cathepsin G (Figure 4k)^72^. In summary, these data highlight an overall pro-inflammatory and potentially tissue-damaging phenotype of peripheral blood monocytes in COVID-19 pneumonia.

### Robust early upregulation of ISG signatures in a large ambulatory SARS-CoV2 infected cohort

Using a cohort of high-risk patients, we detected a distinct immune profile in non-pneumonic SARS-CoV-2 infection, which was most prominently characterized by a global ISG response across different peripheral immune cell subsets, a lymphocyte shift from cytotoxic to immune-regulatory, and an anti-inflammatory monocyte signature. Longitudinal analysis most notably highlighted an early global ISG response to be decisive for a non-pneumonic disease course, whereas the other identified immune trajectories developed over time. Because of the crucial patho-mechanistic and potentially therapeutic implications, we sought to verify our findings by focusing on ISG expression at an early timepoint upon infection. To again concentrate on protective immunity, we chose a prospective cohort of ambulatory patients, which prospectively enrolled SARS-CoV 2 PCR-positive individuals, who were not hospitalized due to paucity of symptoms (validation cohort: KoCo19-Immu, see Methods)^73, 74^. We included PBMC samples early after RT-PCR-positivity (day 4) and after convalescence (day 60) (n=39). Two additional cross-sectional cohorts of age-matched SARS-CoV-2 negative patients (n=9) and a second hospitalized cohort with COVID-19 pneumonia (n=7) were also analyzed for reference. We performed subset bulk RNA-sequencing of sorted major immune cell subsets to allow for a deep transcriptomic coverage (see Methods).

In line with the exploratory cohort, ambulatory SARS-CoV-2 infected patients with oligo-to asymptomatic disease showed an early upregulated ISG signature across major PBMC subsets at day 4 compared to after convalescence at day 60 after infection (Figure 5a). Early in the disease course in comparison to convalescent patients from the same cohort, 33 ISGs in monocytes were upregulated and 6 downregulated. (Figure 5b). Corresponding to our exploratory cohort, *IFI44, IFITM3, IFI44L, MX1* and other ISGs were significantly upregulated in monocytes in ambulatory cases (Figure 5b). Similarly, CD4^+^ T cells showed an upregulation of *IFI44, IFI44L, Ly6E and MX1*, as a few examples of 44 up- and 11 downregulated ISGs in CD4^+^ T cells (Figure 5c). Transcriptional analysis of NK cells revealed upregulation of *XAF1, Ly6E, IFI6 and IFITM3*, with 36 ISGs up- and 15 downregulated altogether in NK cells (Suppl Fig 7a-b). Finally, we computed an ISG score using the same interferons as for the exploratory cohort (Fig 5d). The ISG score of early disease ambulatory patients was significantly increased compared to convalescent patients, a very similar finding to the ISG score derived from the exploratory cohort scRNA Seq data (Fig 2k). A similar immune signature emerged upon comparison of day 4 samples with SARS-CoV-2 negative, age-matched controls (Suppl. Fig. 7c).

**Fig 5.**
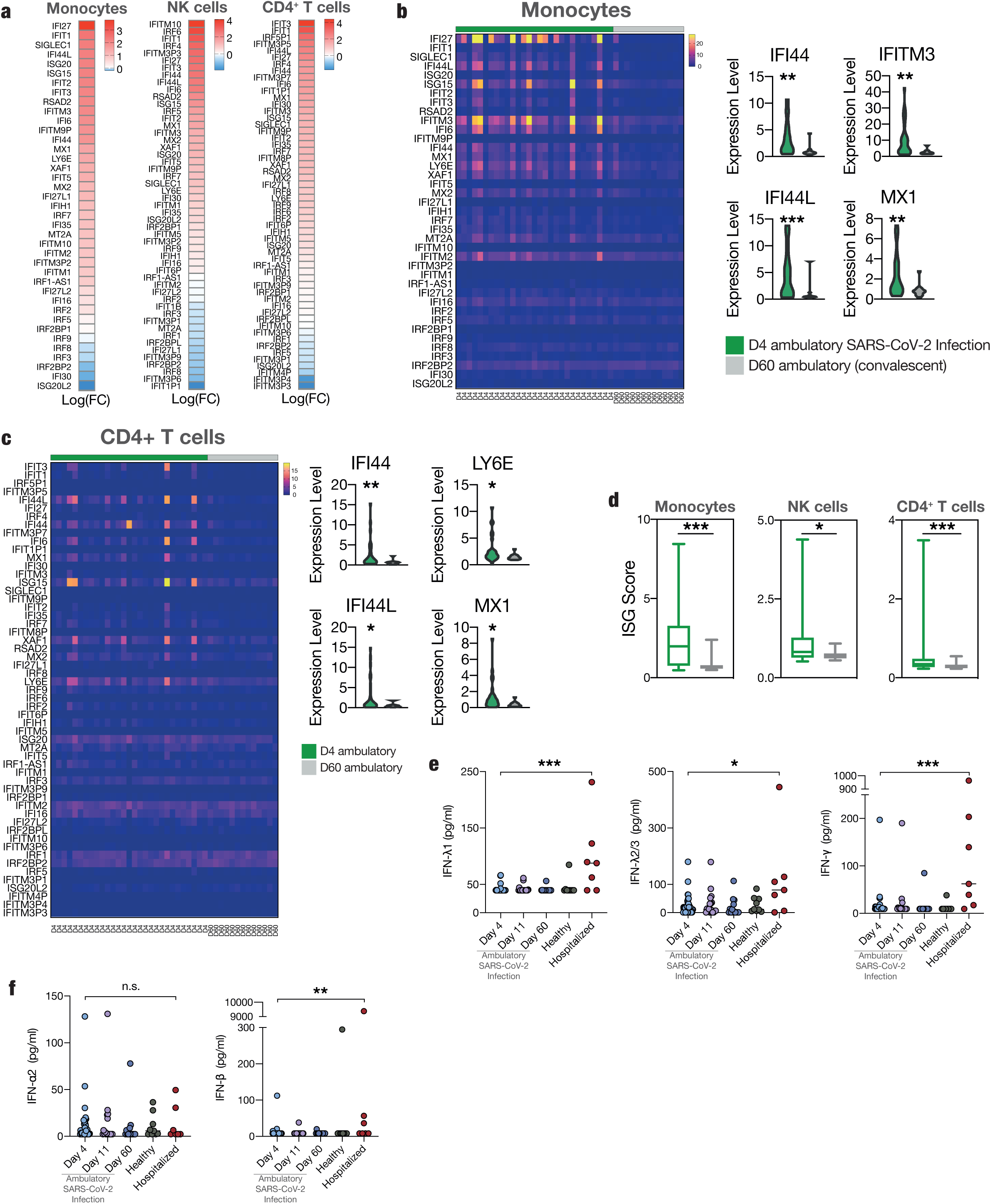
Differential cytokine regulation and robust early upregulation of ISGs in a large ambulatory SARS-CoV-2 infected cohort. **a:** Heat maps of differentially expressed interferon stimulated genes in leukocyte subsets (monocytes, NK cells, CD4^+^ T cells) of day 4 ambulatory compared to day 60 (convalescent) COVID-19. Monocytes: n=33 upregulated, n=6 downregulated. NK cells: n=26 upregulated, n=15 downregulated CD4^+^ T cells: n=44 upregulated, n=7 downregulated. **b-c:** Heat maps and violin plots of differentially expressed interferon stimulated genes in monocytes and CD4^+^ T cells of day 4 ambulatory compared to day 60 (convalescent) COVID-19. Individual ISG expressions of exemplary ISGs. **d:** Computed ISG scores for monocytes, NK cells and CD4^+^ T cells. **c-e**: Mann-Whitney U tests. n=29 d4, n=13 d60. **e-f:** Measurements of IFN-λ1, IFN-λ2/3, IFN-γ **(e)**, IFN-α2, IFN-β **(f)** in plasma samples. D4, 11 and 60 longitudinal ambulatory COVID-19 samples. n=9 controls, n=40 d4, n=18 d11, n=14 d60, n=7 hospitalized COVID-19. Mann-Whitney U tests between d4 and all other groups. Non-significant results not shown, besides for hospitalized. Line denotes median. *p<0.05, **p<0.01, ***p<0.001.

To test if our findings are consistent, we further compared ambulatory SARS-CoV-2 infection with another cohort of hospitalized COVID-19. ISG expression in monocytes and NK-cells of day 4 ambulatory SARS-CoV-2 infected individuals compared to hospitalized patients with severe symptoms and consistent lung involvement again yielded similar results. Despite strong differences in symptom severity, we observed the same trend: In monocytes from ambulatory, infected patients at an early disease stage, 28 ISG were upregulated and 16 downregulated in comparison to hospitalized patients (Suppl. Fig. 7d). Additionally, 25 ISGs were up- and 15 downregulated when comparing NK cells of ambulatory to hospitalized patients (Suppl. Fig. 7e).

In summary, a large, prospective, ambulatory cohort of SARS-CoV-2 infected patients validated a strong early interferon response to characterize a disease course without substantial lung involvement.

### Cytokine profiling reveals differences in type II and III interferon plasma levels

Next, we asked whether the observed ISG signature in circulating immune cells of SARS-CoV-2 infected ambulatory patients (validation cohort) was due to elevated systemic interferon levels. We deployed multiplex cytokine profiling to longitudinally map circulating plasma interferons at day 4, 11 and convalesce (day 60) of ambulatory (n=40) compared to hospitalized COVID-19 (n=7) and non-COVID-19 control patients (n=9).

Interestingly, in contrast to identified ISG responses, non-hospitalized mild cases at early timepoints of infection (d4) had a significantly lower level of interferon type II and type III (IFN-λ1, IFN-λ2/3 and IFN-γ) compared to hospitalized COVID-19 patients (Figure 5e). However, there was no clear difference between early mild and hospitalized SARS-CoV-2 infection for interferon type I plasma levels, with no difference in circulating IFN-α2 and only slightly higher IFN-β in hospitalized patients (Figure 5f). There was no difference between d4 and any other timepoints as well as to healthy controls. In summary, despite overt differences in ISGs, cytokine profiling provided no evidence for differences in circulating IFN levels as an explanation for the observed ISG signature across immune cell subsets in non-pneumonic SARS-CoV-2 infected patients. Indeed, the ISG scores of immune cells of hospitalized and day 4 ambulatory patients and their respective plasma interferon levels did not correlate, with no significant correlations across any cell or interferon type (Suppl Fig 7f). This might indicate either a local ISG induction in the upper airways, independent of plasma-interferons, or interferon-independent inductions of ISG expression^75^. Plasma-interferon independent changes in cellular ISGs might explain the conflicting data currently existing on interferon responses in SARS-CoV-2 infection^13, 18, 19, 76^.

## Discussion

The heterogeneity of responses to SARS-CoV-2 infection, ranging from non-pneumonic courses to acute respiratory distress syndrome (ARDS), might hold promise for immune system modulation as a therapeutic approach. Even the majority of patients with high-risk characteristics are able to control the virus in the upper airways^77^. Here, we specifically analyzed immune responses that enable early viral containment without triggering organ damage by focusing on non-pneumonic compared to pneumonic SARS-CoV-2 infection. We utilized an at-risk cohort analyzed with state-of-the-art multi-omics assays to generate hypotheses, which we then validated in a larger prospective validation cohort of ambulatory infected patients. ^48, 63^ Most importantly, we dissected the features defining a successful immune response against SARS-CoV-2 in non-pneumonic and ambulatory infected individuals.

A global upregulation of ISG responses in non-pneumonic SARS-CoV-2 infection were the most prominent changes observed across major peripheral blood immune cell populations. These ISG signatures were most distinctly upregulated at early disease timepoints, which hints at a causative protective function of these pathways. We confirmed these findings in a prospective ambulatory validation study (validation cohort). Interestingly, these changes in ISG signature were not accompanied by increased plasma interferon levels, and cellular ISGs did not correlate with plasma interferon levels. In addition, we found that non-pneumonic SARS-CoV-2 infection was associated with immunomodulatory T cell function with reduced cytotoxicity and a pro-resolving EGFR signature in monocytes. In combination, these trajectories are likely to limit uncontrolled cytokine release and inflammation without simultaneously interfering with a locally and temporally restricted innate and adaptive (mucosal) immune response^67, 78^.

In contrast, pneumonic patients showed a reduced early ISG response, but displayed an upregulation of monocyte IL6R, S100A8/A9, and an increased cytotoxicity of T cells and NK. This shift to a proinflammatory phenotype after a failed ISG response might represent a compensatory response, triggering tissue damage and explain immunopathology and cytokine release associated with poor outcome^79^.

Conflicting data exist on IFN signaling in COVID-19 depending on (1) patient collective, (2) sampling time points and (3) disease state^13, 18, 19, 76^. It still remains unclear which role plasma interferons and interferon stimulated intracellular pathways play for an uncomplicated disease course. Our work strongly supports the hypothesis of an early pronounced ISG response as key for limiting viral spread at early timepoints, whereas a delayed/suppressed response leads to viral dissemination and an immunopathogenic global inflammatory response^80^. Interestingly, cytokine profiling excluded concomitant increases in circulating IFN I and III levels in oligo- and asymptomatic individuals, which poses the question how ISG upregulation is mediated in these cases. Possibly, IFN produced locally in first and second tier immune responses of upper airway and lymphatic tissues primes circulating immune cells^81^. Also, non-canonical ISG upregulation independent of classical IFN receptor-JAK-STAT signaling has been described in viral infection^82, 83^. Alternatively, we cannot exclude a very early, transient IFN release not captured by our sampling time points. Further research is needed to pin down mechanisms of a systemic anti-viral immune state and downstream effector pathways in non-pneumonic SARS-CoV-2 infection.

Our data from non-pneumonic patients bridge the gap to recent insights into deranged IFN responses in severe COVID-19, which are at least in part caused by inborn or acquired defects in interferon signaling^12, 15, 16, 84^. Local upregulation of ISG-signatures in upper airway epithelium has been associated with successful virus clearance in influenza^85^. This might hold important therapeutic and diagnostic implications. Measuring systemic ISG response in early disease might help to differentiate favorable outcomes from severe disease courses, but this needs further evaluation in a prospective study.

Our data show that therapeutic induction of ISGs in immune cells might be a treatment option in early disease. So far, data on interferon-based therapy in COVID-19 is inconclusive, but a study utilizing inhaled nebulized interferon beta-1 in non-severe COVID-19 showed promising results^86, 87^. Intriguingly, retrospective analysis of a cohort trial with IFN-alpha2 indicated that early interferon treatment might be beneficial regarding clinical outcomes, with harmful effects late in the disease^88^. However, side-effects and the abovementioned autoantibodies as well as signaling mutations in patients susceptible to SARS-CoV-2 might limit the use of interferon-alpha or beta^15, 16, 89^.

In summary, our data provide important implications for therapy and disease course prediction in SARS-CoV-2 infection. By measuring ISG responses in peripheral immune cells, patients could potentially be segregated in ARDS high/low risk cohorts. Treatment with interferons early in disease might be able to tip the inflammatory response towards local pathogen control, possibly preventing severe COVID-19 and its potentially fatal consequences.

## Acknowledgments

The authors thank the patients and their families for blood donations and for their participation in the CORKUM registry. We would like to thank all CORKUM investigators and staff.

## Authorship contributions

Initiation: K.P., K.S., L.N.; Conceptualization: K.P., A.L., L.N.; Methodology: K.P., A.L., R.K., L.N.; Investigation: K.P., A.L., R.K., S.B., A.J., O.P., V.P., L.E.W., C.G., M.K., M.M., J.C.H., C.S., T.E., F.D., N.K., A.Li., K.S., L.T., L.N., M.J.; Resources: S.M., K.S., Ch.S., J.C.H., W.E., R.Z., I.K., C.Ge., P.M., N.H., M.H.; Formal analysis: K.P., A.L., R.K., S.B., M.J., O.P., P.M., L.N.; Visualization: A.L., M.J., O.P.; Supervision: K.P., K.S., L.N. R.Z.; Project administration: K.P., K.S., L.N.; Funding Acquisition: K.P., S.M., K.S., L.N.; Writing original draft: K.P., A.L., R.K., L.N.; Editing: All authors.

## Data availability

Data available upon reasonable request from the authors.

