## Supplementary material for "Protective immune trajectories in early viral containment of non-pneumonic SARS-CoV-2 infection": Suppl. Data and Figures

Suppl. Methods and Figure Legends.

Pekayvaz et al.

### Methods

#### Ethics

In accordance with the Declaration of Helsinki, and with the approval of the Ethics Committee of Ludwig-Maximilian-University Munich, informed consent of the patients or their guardians was obtained. COVID-19 patients are part of the COVID-19 Registry of the LMU University Hospital Munich (CORKUM, WHO trial ID DRKS00021225). Pseudonymized data was used for analysis, the CORKUM and KocoImmu studies were approved by the ethics committee of LMU Munich (No: 20-245 & No: 20-371 respectively).

#### Cohorts

A total of 126 samples from 72 patients were included in this study. Two independent cohorts were used, an exploratory cohort (scRNA-Seq, flow cytometry, plasma proteomics) and a confirmation cohort (leukocyte subset in-depth RNA-Seq, cytokine assay).

*Exploratory cohort:* For our exploratory cohort, a subset of patients without any pulmonary involvement were included as a model immune response to SARS-CoV-2. In total, 14 subjects were included in our exploratory cohort (n=11 patients with positive SARS-CoV-2 RT-PCR and n=3 non-COVID-19 control subjects). COVID-19 patients were sampled longitudinally, and three time points were included: first as soon as possible after positive RT-PCR or symptom onset, second in between the first and third time point, third after seroconversion, negative RT-PCR or symptom improvement. 14 patients were included for flow cytometric analysis, 12 patients were included into single cell RNA-Seq assays. Patients with severe pre-existing kidney or liver dysfunction, severe autoimmune diseases, chronic infection, patients requiring ECMO therapy, with a known coinfection with Influenza or Respiratory Syncytial Virus (RSV) were excluded. COVID-19 patients were divided into patients without any pulmonary symptoms or radiological infiltrates and patients with confirmed COVID-19 associated pneumonia. Furthermore, control subjects without COVID-19 were included. Average number of risk factors was calculated based on individual risk factor sum. Risk factors were medical or physiological conditions associated with severe COVID-19: Age>60yrs, arterial hypertension, cardiovascular disease, chronic respiratory disease, diabetes mellitus, and male gender[90].

*Confirmation cohort:* The confirmation cohort consisted of a total of 58 patients. Of these, n=42 were SARS-CoV-2 positive, non-hospitalized individuals. These 42 patients participated in the longitudinal KoCo19-Immu cohort, which enrolled SARS CoV-2 infected individuals shortly after PCR confirmation. Comprehensive longitudinal blood sampling was performed by household visits of field teams. The ambulatory SARS CoV-2 infected individuals were analyzed at three time points after initial RT- PCR confirmed COVID- 19 infection: first at day 4 after RT-PCR, second at day 11 and third at day 60 after positive RT-PCR. The median day for the first timepoint was on day 6 after symptom onset (IQR: 5 to 9.75 days), the median for the second was on day 15 [IQR: 11.75 – 17.25] and for the last visit on day 68 [IQR: 63.25 – 83.25] after symptom onset. A subset of n=40 longitudinally sampled, ambulatory patients were used for plasma cytokine analysis, with n=40 d4, n=18 d11, n=14 d60 samples measured. Another subset of n=39 ambulatory patients were used for subset RNA sequencing, with n=39 d4, n=13 d60 samples used. In addition to these 42 ambulatory patients, a reference cohort of SARS-CoV-2 (n=9) negative individuals (female: 78%, median age: 27) and hospitalized COVID-19 (n=7) were recruited (female: 29%, median age: 82). The confirmation cohort was sampled and processed completely independently from the exploratory cohort to reduce any systemic bias. The KoCo19-Immu-study is conducted under the framework of the prospective population based Koco19 cohort [72, 73].

### Peripheral blood mononuclear cell (PBMC), plasma collection and storage

For plasma isolation, heparin anticoagulated blood was centrifuged for 10 minutes at 650 g and the plasma layer was collected. Red and white blood cells were diluted at a ratio of ~ 1:2 - 1:3 with PBS. 35ml of PBS-Blood suspension was slowly transferred on top of a 15ml Pancoll (PAN Biotech; Cat.: P04-60500) solution to create a blood layer and subsequently centrifuged for 40minutes at 700 g with the slowest acceleration and deceleration program. The buffy coat was isolated and washed twice with PBS (at 600 g for 7:30 minutes), resuspended in RPMI, counted, washed again with PBS and then frozen at a concentration of $1x10^7$ cells/ml in freezing medium (90% FCS + 10% DMSO). For the validation cohort CPDA (Citrat-Phosphat-Dextrose-Adenin) blood was centrifuged at 1285 g for 10 minutes. Plasma was removed and stored subsequently and twice the amount of PBS was added to the cell pellet. PBS / Blood suspension was added to Leucosept tubes (Greiner) with a Ficoll-Paque at a 1:2 ratio and centrifuged at 800 g, the PBMC fraction was isolated subsequently. Cells were slowly frozen in a Mister Frosty for 24 h at -80°C, and then transferred to a liquid nitrogen tank.

### PBMC processing and preparation for flow cytometry and FACS/Sorting

PBMC vials were thawed at room-temperature for 10 minutes and transferred to 5 ml PBS with 1% bovine serum albumin (BSA). Cell suspension was centrifuged at 400 x g for 15 minutes at 4 °C. Hereafter, cells were resuspended in PBS with 1% BSA and stained on ice with respective antibody master-mix panels for subsequent flow cytometry analysis and FACS-sorting.

### FACS/Sorting for single-cell RNA-seq

Cells were stained with SYTOX™ Red (Cat No. 1936399, Invitrogen) prior to sort. $CD45^+$ living singlets were FACS/sorted and centrifuged at 400 x g for 10 minutes at 4°C. The cell concentration was adjusted to 800 cells / µl in PBS.

### Single-cell RNA-seq

The Chromium Next GEM Single Cell 3’ Reagent Kit with Feature Barcoding technology (CG000206 Rev D) was used. For feature Barcoding, patient samples were tagged by TotalSeq™ anti-human Hashtag Antibodies (B0251, A0252, A0253). Hashtag antibodies were included into the antibody panel for FACS/Sorting. 3 patient samples from one timepoint and group were pooled per library. In brief, according to the manufacturer’s instructions, first Gel Beads-in-emulsion (GEMs) were generated, reverse transcribed, cleaned up and cDNA was amplified. After cDNA generation and amplification, cDNA was quality controlled and quantified. Subsequently, the 3’ gene expression library and the cell surface protein library was constructed. Sequencing was performed with Illumina NovaSeq (library preparation and sequencing was performed by IMGM laboratories).

### Single-cell RNA-seq data processing

The raw reads were obtained from the sequencing facility. A total of 12 samples were processed using cellranger 4.0.0 with the 10X human reference data GRCh38 2020A (https://support.10xgenomics.com/single-cell-gene-expression/software/release-notes/build#GRCh38_2020A).

The resulting filtered count matrices were loaded using Seurat 3.2.1[91], filtered (nFeature_RNA > 200, nFeature_RNA < 6000, nCount_RNA > 1000, percent.rp < 40, percent.mt < 15), normalized (SCTransform [3]) and integrated (according to Seurat's SCTransform integration vignette) into one combined Seurat object. On the combined Seurat object PCA was performed with default parameters and subsequent calculation of the UMAP embedding (using the first 30 PCs). FindNeighbors was called with default parameters. Seurat's FindClusters was called with a resolution of 0.5. Gene expression was quantified for each cluster's marker genes, which were determined by Seurat's FindMarkers function using the t-test. Using the cluster marker genes and gene expression results cell types were predicted using the scRNA-seq cell type prediction [92]on PanglaoDB's marker genes [93] and restricted to an 'Immune system' context. After manual curation of the predictions by the experimentalists (cluster 1 NK -> T, 4 Mono -> T, 13 NK-> T, 15 Mac -> Mono), a cell type was assigned to each cluster. It should be noted that for all these manual curations, the finally selected cell type was the second highest scoring prediction. Gene set enrichment analysis (overrepresentation analysis) on Gene Ontology (Biological Process aspect) [94] was performed using clusterProfiler[95]. The interferon score is calculated as mean over the normalized expression values of the interferon genes (*IFI6, IFI27, IFI44, IFI44L, IFIT1, IFIT2, IFIT3, IFITM1, IFITM3, ISG15, LY6E, MT2A, MX1, RSAD2, SIGLEC1*) for each cell. Significance between groups is calculated using the t-test with 'ggpubr' (https://CRAN.R-project.org/package=ggpubr) stat_compare_means function. The full analysis script is available from GitHub at https://github.com/mjoppich/covidSC in the analysis_final_analysis.Rmd script. This repository also contains all scripts used for set enrichment. Functions for custom plots are included in the main analysis script.

**Peripheral blood mononuclear cell (PBMC) processing for subset RNA-seq**

PBMCs were thawed at 37 °C for 5 minutes, added to 5ml PBS with 1% BSA, centrifuged at 350xg for 10 minutes at 4°C. The supernatant was removed and resuspended in 150 µl PBS with 0,5% BSA. 50 µl was added to 50 µl of panel 1 and panel 2 respectively and incubated on ice for 15 minutes. Probes were filtered through a 50 µm mesh and 0,1 µl of Sytox Red (Cat No. 1936399, Invitrogen) was added to the probe prior to Sort. $CD4^+$, $CD8^+$ T cells, NK cells and monocytes were sorted. 500-1000 cells were sorted into 100ul Buffer RLT Plus (+1% Mercapto-Ethanol).

| Sort Panel 1 | | |
|---|---|---|
| **Color** | **Antigen** | **Company & Cat #** |
| PE | CD3 CD20 | BioLegend #300308, #302306 |
| FITC | CD14 | BD Biosciences #557153 |
| PE-Cy7 | CD16 | BD Biosciences #557744 |
| APC-Cy7 | CD56 | BioLegend #362512 |

| Sort Panel 2 | | |
|---|---|---|
| **Color** | **Antigen** | **Company & Cat #** |
| PE | CD3 | BioLegend #300308 |
| PE-Cy7 | CD4 | BioLegend #357410 |
| AF488 | CD8a | BioLegend #301024 |
| APC-Cy7 | CD19 | BioLegend #363010 |

**Library preparation and subset RNA-seq**

RNA-sequencing was performed using prime-seq. A step-by step protocol can be found on protocols.io (dx.doi.org/10.17504/protocols.io.s9veh66).

Briefly, of the 1000 cells that were sorted in the lysis buffer, 50 µL (500 cells) were used to prepare RNA-seq libraries. The lysate was treated with Proteinase K (AM2548, Life Technologies), isolated with cleanup beads (GE65152105050250, Sigma-Aldrich) (2:1 beads/sample ratio), and then DNase I (EN0521, Thermo Fisher) digested. The RNA was then reverse transcribed with 30 units of Maxima H- enzyme (EP0753, Thermo Fisher), 1x Maxima H- Buffer (EP0753, Thermo Fisher), 1 mM each dNTPs (R0186, Thermo Fisher), 1 µM template-switching oligo (IDT), 1 µM barcoded oligo-dT primers (IDT) in a 10 µL reaction volume at 42 °C for 90 minutes. The samples belonging to the same tissue were then pooled and cleaned using cleanup beads (1:1 beads/sample ratio), resulting in 4 pools, one for monocytes, NK cells, CD4 T cells and CD8 T cells. Following cleanup, remaining primers were digested with

Exonuclease I (M0293L, NEB) at 37 °C for 20 minutes followed by 80 °C for 10 minutes. The Exonuclease I digested samples were then again cleaned using cleanup beads (1:1 beads/sample ratio).

Second strand synthesis and pre-amplification was performed using 1X KAPA HS Ready Mix (07958935001, Roche) and 0.6 µM SINGV6 primer (IDT) in a 50 µL reaction. The PCR was cycled as follows: 98 °C for 3 minutes; 12 cycles of 98 °C for 15 s, 65 °C for 30 s, 72 °C for 4 minutes; and 72 °C for 10 minutes. The samples were then cleaned using cleanup beads (0.8:1 beads/sample ratio) and then eluted in 10 µL of DNase/RNase-Free Distilled Water (10977-049, ThermoFisher). The Quant-iT PicoGreen dsDNA Assay Kit (P7581, Thermo Fisher) was used to quantify the amount of cDNA present, and the High-Sensitivity DNA Kit (5067-4627, Agilent) was used to qualify the size distribution.

Following QC, 2.5 µL of cDNA (2.25 - 5.75 ng) from each sample was used to make libraries with the NEBNext Ultra II FS Library Preparation Kit (E6177S, NEB), primarily following manufacturer's instructions but with a five-fold lower reaction volume. Fragmentation was carried out using the supplied Enzyme Mix and Reaction buffer in a 6 µL reaction. The adapters were ligated using the supplied Ligation Master Mix, Ligation Enhancer, and a custom prime-seq Adapter (1.5 µM, IDT) in a reaction volume of 12.7 µL. Following ligation, the samples were double-size selected using SPRI-select Beads (B23317, Beckman Coulter), with 0.5 and 0.7 ratios. The samples were then amplified using a library PCR using Q5 Master Mix (M0544L, NEB), 1 µL i7 Index primer (Sigma-Aldrich), and 1 µL i5 Index primer (IDT) using the following setup: 98 °C for 30 s; 13 cycles of 98 °C for 10 s, 65 °C for 1 m 15 s, 65 °C for 5 m; and 65 °C for 4 m. A final double-size selection was performed as before using SPRI-select Beads.

After checking the concentration and quality using a high-sensitivity DNA chip (Agilent Bioanalyzer), the libraries were 150 bp paired-end sequenced on a S4 flow cell of a NovaSeq *** (Ilumina).

The data was initially checked using fastqc (version 0.11.8[96]). Cutadapt (version 1.12[97]) was then used to remove any regions on the 3' end of the read where the sequence read into the polyA tail. Following processing of the data, the zUMIs pipeline (version 2.9.4d, Parekh et al., 2018) was used to filter the data, using a phred threshold of 20 for 4 bases for both the UMI and BC, map the reads to the human genome (GRCh38) with the Gencode annotation (v35) using STAR (version 2.7.3a), and count the reads using RSubread (version 1.32.4)[98, 99].

**Subset RNA-seq data analysis**

Count matrices for each cell type and two counting methods were received. The count matrices with exon counts (exon) and combined intron+exon counts (inex) were extracted for each cell type using the 'all' slot (non-downsampled) from zUMI's data object. The samples and conditions were annotated with the sample names. Both count matrices were processed analogously and differentially expressed genes were determined with DESeq2[100]. For the subsequent data analysis, the exon-count results are used with an adjusted (Benjamini-Hochberg) p-value cut off at 0.05. Gene set enrichment analysis (overrepresentation analysis) on Gene Ontology (Biological Process aspect) was performed using clusterProfiler[94, 95].

**Flow cytometry**

After sample preparations for scRNA-seq and flow cytometry, 50µl of the cell suspension was incubated for 20min on ice with 50µl of the respective antibody panel, at 1:100 dilution for each antibody. After centrifugation at 400g for 7min and resuspension in 200µl 1% BSA with PBS, 0.2µl of SYTOX™ Blue (Cat No. 2192317, Invitrogen) was added for live/dead staining. To avoid batch effects all samples were measured during one experimental run.

Measurements were done on a BD LSRFortessa Flow Cytometer. Analysis was done using FlowJo Software (FlowJo v. 10.6.1, BD). The gating strategy used is shown in Suppl Fig 1d. After a common gating strategy, each cell population/antibody panel was gated separately. n=7 pneumonic, n=4 non-pneumonic COVID-19 and n=3 control patients were used. All three different time points for the COVID samples were measured. For t-SNE and FlowSOM analysis, cell populations were downsampled using the downsample v3 plugin for FlowJo to 2000 cells per time point and sample where possible and subsequently concatenated. FlowSOM clustering with the FlowSOM v2.5 plugin[101] was performed on the concatenated file with n=5 clusters for the NK cell panel and n=8 clusters for both T cell panels and the monocyte panel. The FlowSOM output used is the minimum spanning tree, heatmap of surface maker expression by FlowSOM cluster and the the percentage of cells in each cluster per group and time point. For tSNE and FlowSOM only surface markers that were not previously used for positive cell identification were used.

| NK Cell Panel | | |
|---|---|---|
| **Color** | **Antigen** | **Company & Cat #** |
| BV650 | CD45 | BioLegend #304044 |
| FITC | CD94 | BioLegend #305504 |
| PE | CD3 | BioLegend #300308 |
| PE | CD20 | BioLegend #302306 |
| PERCP-Cy5.5 | CD44 (RAT) | BioLegend #103032 |
| APC | CD160 | BioLegend #341208 |
| APC-Cy7 | CX3CR1 (RAT) | BioLegend #341616 |
| BV 605 | CD161 | BioLegend #339916 |
| BV785 | CD62L | BioLegend #304830 |
| BV510 | CD16 | BioLegend #302048 |
| BV 711 | CD56 | BioLegend #362542 |
| AF 700 | CD18 | BioLegend #302124 |
| PE-Dazzle | CD52 | BioLegend #316014 |
| PE-Cy7 | CD9 | BioLegend #312116 |

| Monocyte Panel | | |
|---|---|---|
| **Color** | **Antigen** | **Company & Cat #** |
| BV650 | CD45 | BioLegend #304044 |
| FITC | CD14 | BD Biosciences #557153 |
| PE | CD3 | BioLegend #300308 |
| PE | CD20 | BioLegend #302306 |
| PE | CD56 | BioLegend #304606 |
| PERCP-Cy5.5 | CD1c | BioLegend #331514 |
| APC | CD45RA | BioLegend #304112 |
| APC-Cy7 | CD88 | BioLegend #344316 |
| BV 605 | HLA-DR | BioLegend #307640 |
| BV785 | CD123 | BioLegend #306032 |
| BV510 | CD16 | BioLegend #302048 |
| BV 711 | CD2 | BioLegend #300232 |
| AF 700 | CD5 | BioLegend #364026 |

| T Cell Panel 1 | | |
|---|---|---|
| **Color** | **Antigen** | **Company & Cat #** |
| BV650 | CD8 | BioLegend #344730 |
| FITC | CD25 | BioLegend #302604 |
| PE | CCR10 | BioLegend #341504 |
| PERCP-Cy5.5 | CD152 (CTLA-4) | BioLegend #369608 |
| APC | CD45RA | BioLegend #304112 |
| APC-Cy7 | CD127 (IL7Ra) | BioLegend #135040 |
| BV605 | CD4 | BioLegend #317438 |
| BV785 | CXCR3 | BioLegend #353738 |
| BV510 | CD3 | BioLegend #317332 |
| BV711 | CD196 (CCR6) | BioLegend #353436 |

| | | |
|---|---|---|
| AF700 | CD197 | BioLegend #353244 |
| PE-Dazzle | CCR4 | BioLegend #359420 |
| PE-Cy7 | CD279 (PD-1) | BioLegend #621616 |

| T Cell Panel 2 | | |
|---|---|---|
| **Color** | **Antigen** | **Company & Cat #** |
| BV650 | CD8 | BioLegend #344730 |
| FITC | CD25 | BioLegend #302604 |
| PE | CD95 | BioLegend #305608 |
| PERCP-Cy5.5 | CD44 (RAT) | BioLegend #103032 |
| APC | CD45RA | BioLegend #304112 |
| APC-Cy7 | CX3CR1 (RAT) | BioLegend #341616 |
| BV 605 | CD4 | BioLegend #317438 |
| BV785 | CD62L | BioLegend #304830 |
| BV510 | CD3 | BioLegend #317332 |
| BV 711 | CD11b | BioLegend #301344 |
| AF 700 | CD197 | BioLegend #353244 |
| PE-Dazzle | CD45RO | BioLegend #304248 |
| PE-Cy7 | CD27 | BioLegend #356412 |

**Plasma sample preparation for mass spectrometry and analysis**

Samples were boiled in SDS-buffer and subjected to SP3-based cleanup and tryptic digest as described[102]. The method was adapted in-house to work on an AssayMAP Bravo Protein Sample Prep Platform (Agilent). A spectral library was generated by measuring 52 fractions of a high-pH fractionation using a uniform peptide mix of all samples in data-dependent mode on a Q Exactive HF-X orbitrap mass spectrometer (Thermo Fisher Scientific). For data-independent acquisition, each individual sample was injected in at least technical duplicates and measured in data-independent mode with variable isolation window sizes. Spectral library generation as well as DIA analysis was performed in Spectronaut (version 14.3) using the Qvalue sparse setting with a q-value of 0.01 in at least one raw-file using the top3 precursors for quantitation without applying imputation[103]. Further downstream analysis was performed in R. Technical replicates were collapsed into samples by using the average Quantity value. For significance calling on protein level, ratios to the average no-COVID control were calculated, scaled by median-MAD normalization and a one-sample moderated t-test (limma package[104]) was applied among experimental groups. Proteins with a Benjamini-Hochberg adjusted p-value of 0.05 (5% FDR) were considered as significant. Top abundant proteins were subjected to STRING analysis using experiments and databases as interaction sources [105]. For single-sample gene-set enrichment analysis (ssGSEA;[106]), median-MAD scaled log-fold change values of group-wise comparisons were calculated using the mean protein intensity values among each group. Significance calling of pathways was applied on the adjusted p-values from the ssGSEA results.

**Interferon plasma assay**

For longitudinal assessment of type 1/2/3 interferons plasma levels, plasma samples of control (n=9), non-pneumonic SARS-CoV-2 infected patients (day 4 n=40, day 11 n=18, day 60 n=14) and hospitalized patients suffering from COVID-19 pneumonia (n=7) were analyzed using a commercial multiplex bead-based assay (LEGENDplex™ Human Type 1/2/3 Interferon Panel, V-bottom plate, Biolegend #740396) according to the manufacturer's instructions. In brief, frozen plasma samples were thawed to room temperature (RT) and centrifuged (1000 rpm, 1 min). Samples were incubated with assay reagents in the dark for 2 hours on a plate shaker (RT, 600 rpm). After addition of the detection antibody mix, samples were incubated for another hour (RT, 600 rpm), followed by addition of Streptavidin-PE (30 min, RT, 600 rpm) and an on-plate washing step. Finally, samples were measured in technical duplicates using a BD LSRFortessa flow cytometer. Interferon plasma levels were calculated using LEGENDplex™ Data Analysis Software Suite QOGNIT.

#### Data analysis and statistics

Excel (Microsoft), Prism (GraphPad) and R with package *corrplot* were used for data analysis. Adobe Illustrator was used to assemble the graphical illustrations. Results are shown and mentioned in the text as mean ± s.e.m., unless otherwise indicated. For direct comparisons between two groups, unpaired, two-tailed Student's t-tests or Mann-Whitney U tests were used. For changes across clusters in flow cytometric measurements mixed-effects model analysis was used. If the mixed-effects model analysis was significant, a line denotes the significance. Post-hoc Sidak's multiple comparisons test was used for individual significant differences. P values of ≤0.05 are considered significant and denoted with *, ≤0.01 with ** and ≤0.001 with ***. Individual patients or samples are represented as dots, unless otherwise indicated. Error bars are standard error of the mean (s.e.m.) unless otherwise indicated.

SUPPLEMENTARY TABLE 1: Additional clinical characteristics of exploratory patient cohort

| **Cohort:** | **Control** | **non-pneumonic COVID-19** | **pneumonic COVID-19** | **t test / Fisher's exact test non-pneumonic vs. pneumonic** |
|---|---|---|---|---|
| Patient count: | 3 | 4 | 7 | |
| Age, Median Year [interquartile range [IQR]] | 74 [69-74] | 72 [41-88] | 61 [37-75] | n.s. |
| Male, n [%] | 2 [66] | 4 [100] | 4 [57] | n.s. |
| **COVID-19 risk factors** | | | | |
| Cardiovascular disease, n [%] | 3 [100] | 3 [75] | 2 [29] | n.s. |
| Arterial hypertension, n [%] | 3 [100] | 3 [75] | 3 [43] | n.s. |
| Diabetes mellitus, n [%] | 2 [66] | 2 [50] | 3 [43] | n.s. |
| Asthma / COPD / OSAS, n [%] | 1 [33] | 1 [25] | 1 [14] | n.s. |
| Male, n [%] | 2 [66] | 4 [100] | 4 [57] | n.s. |
| Age >60, n [%] | 3 [100] | 2 [50] | 4 [57] | n.s. |
| ≥1 risk factors for severe COVID-19, n [%] | 3 [100] | 4[100] | 7 [100] | n.s. |
| **Pathogens (if tested)** | | | | |
| SARS-CoV-2, n [% positive result] | 0 [0] | 4 [100] | 7 [100] | |
| Other pathogens (Influenza, RSV, etc.), n [% positive result] | 0 [0] | 0 [0] | 0 [0] | |
| **Clinical information at admission** | | | | |
| COVID-19 typical symptoms (New onset fever, cough or dyspnea), n [%] | 0 [0] | 2 [50] | 7 [100] | n.s. |
| **Clinical information during enrollment in the study** | | | | |

| | | | | |
|---|---|---|---|---|
| O2-requirement | 0 [0] | 0 [0] | 6 [86] | * |
| Immunomodulatory treatment / trial enrollment | 0 [0] | 0 [0] | 3 [43] | n.s. |
| **Radiological findings** | | | | |
| Chest CT, n [%] | 3 [100] | 3 [75] | 7 [100] | |
| COVID-19 typical bipulmonary infiltrates, n [%] | 0 [0] | 0 [0] | 7 [100] | ** |
| COVID-19 typical ground glass opacities, n [%] | 0 [0] | 0 [0] | 7 [100] | ** |

*p<0.05,
**p<0.01.

### Suppl. Figure legends.

**Suppl Fig 1 | a:** Schematic of exploratory cohort and sampling time points. **b:** Timeline of the exploratory group normalized by the first sampling time point (day=0). Sampling time points, positive and negative real time polymerase chain reaction (RT-PCR) for SARS-CoV-2, positive and negative COVID-19 in chest tomography/chest x-ray and positive SARS-CoV-2 antibody detection are shown for n=7 pneumonic and n=4 non-pneumonic patients. **c:** Longitudinal blood lymphocyte, neutrophil and monocyte counts at time points 1-3 of n=7 pneumonic COVID-19, n≥3 non-pneumonic COVID-19 patients per time point. No statistical testing performed **d:** Gating strategies for flow-cytometric characterization of PBMCs. Common gating strategy until * is shown in the top row. Further gating strategy is shown for the respective cell population. For dendritic cells, all samples were gated together as a concatenated file. **e:** Dot-plot of cluster defining markers for the scRNA-seq data. **f:** Dot-plot of population defining markers for the scRNA-seq data as shown in Figure 1g.

**Suppl Fig 2 | a:** UMAP representation of the exploratory cohort samples showing the assigned cell populations per sampling time point. **b:** Correlation analyses between plasma proteomic measurement and clinical laboratory measurement of CRP and fibrinogen (FGA). Pearson correlation coefficient shown on top, gray area is 95% confidence interval. **c:** Principle component analysis (PCA) of analysed proteome samples. **d:** Pearson correlation matrix between this proteomic analysis and the proteome from *Shu et al.* Input values are normalized logFC to healthy control samples. **e:** Significantly (adj p<0.05) differentially expressed proteins in plasma samples pooled across all three time points. Log fold change is computed relative to the control protein expression. **f-g:** Volcano plot of differentially expressed proteins of pneumonic (upper) and non-pneumonic (lower) samples per time point compared to controls. Top proteins are annotated. For proteomics: n=4 patients with 12 samples for non-pneumonic and n=6 patients with 18 samples for pneumonic and n=3 samples and patients for controls.

**Suppl Fig 3 | a:** Dot-plot of interferon stimulated gen expression and Percent expressing cells in CD4$^+$ T cells and NK cells per time point in non-pneumonic patients**. b:** Volcano plot of differentially regulated genes in CD4$^+$ T cells, NK cells and monocytes of non-pneumonic compared to control samples. Red annotations are significantly upregulated (adj p val<0.05). Positive fold change signifies higher expression in the non-pneumonic group. Line denotes adj p val<0.05. **c:** GO-BP network analysis of pathways upregulated in CD4$^+$ T cells of non-pneumonic samples compared to pneumonic samples. **d-f:** Violin plots of expression of interferon stimulated in CD4$^+$ T cells, NK cells, C cells and monocytes for pneumonic, non-pneumonic and control samples.

**Suppl Fig 4 | a:** Violin plots of expression of GZMB, LGALS1, S100A8 and S100A9 in NK cells per sampling time point. **b:** GO-BP network analysis of pathways upregulated in NK cells of pneumonic samples compared to non-pneumonic samples. **c:** tSNE of NK cells measured by flow cytometry. FlowSOM assigned clusters are shown in different colors. **d:** Heat map of surface marker expression by FlowSOM group. **e:** FlowSOM self-organizing map of NK cells. **f:** Violin plots of Percentage of NK cells in each FlowSOM cluster per time point. Mixed-effects model analysis. Lines denote significant time effect, post-hoc Sidak's multiple comparisons test for individual significant differences between pneumonic and non-pneumonic samples per time point. n≥3 per time point and group. **g:** tSNE with expression for individual surface makers of NK cells. **h:** GO-BP network analysis of pathways enriched in CD8$^+$ T cells of pneumonic samples compared to non-pneumonic samples. **i:** Violin plots of expression of cytotoxicity related genes in NK cells and CD8$^+$ T cells for pneumonic, non-pneumonic and control samples. **j:** Volcano plots of differentially expressed plasma proteins at TP2-3 of pneumonic samples compared to control samples. Line denotes adj p val<0.05. **k:** String network of top upregulated proteins in pneumonic plasma samples. *p<0.05, **p<0.01, ***p<0.001.

**Suppl Fig 5 | a:** Percentage of CD4$^+$ and CD45RA/RO$^{+/-}$ CD4$^+$ T cells of live PBMCs measured by flow cytometry per sampling time point. Two-sided t-test, n≥3 per time point and group. **b:** tSNE of T cells (T cell panel 1) measured by flow cytometry. FlowSOM assigned clusters are shown in different colors. **c:** Heat map of surface marker expression by FlowSOM group. **d:** Violin plots of Percentage of T cells in each FlowSOM cluster per time point. Mixed-effects model analysis. Lines denote significant time effect, post-hoc Sidak's multiple comparisons test for individual significant differences between pneumonic and non-pneumonic samples per time point. n≥4 per time point and group. **e:** tSNE with expression for individual surface makers of T cells. **f:** FlowSOM self-organizing map of T cells. **g:** tSNE of T cells (T cell panel 2) measured by flow cytometry. FlowSOM assigned clusters are shown in different colors. **h:** Heat map of surface marker expression by FlowSOM group. **i:** Violin plots of Percentage of T cells in each FlowSOM cluster per time point. Mixed-effects model analysis. Lines denote significant time effect, post-hoc Sidak's multiple comparisons test for individual significant differences between pneumonic and non-pneumonic samples per time point. n≥4 per time point and group. **j:** FlowSOM self-organizing map of T cells (T cell panel 2). **k:** tSNE with expression for individual surface makers of T cells (T cell panel 2). All error bars are mean ± s.e.m. unless otherwise noted. *p<0.05, **p<0.01.

**Suppl Fig 6 | a:** Dot-plot of selected gene expression and percent expressing cells in non-pneumonic NK and T cells per time point. **b:** tSNE of monocytes measured by flow cytometry. FlowSOM assigned clusters are shown in different colors. **c:** Violin plots of Percentage of monocytes in each FlowSOM cluster per time point. Mixed-effects model analysis. Lines denote significant time effect, post-hoc Sidak's multiple comparisons test for individual significant differences between pneumonic and non-pneumonic samples per time point. n≥3 per time point and group. **d:** FlowSOM self-organizing map of monocytes **e:** tSNE with expression for individual surface makers of monocytes. *p<0.05, **p<0.01.

**Suppl Fig 7 | a:** Heat map of differentially expressed interferon stimulated genes in NK cells of day 4 ambulatory compared to day 60 (convalescent) SARS-CoV-2 infection. **b:** Individual ISG expressions of exemplary ISGs. Mann-Whitney U tests. n=29 d4, n=13 d60. **c:** Computed ISG scores for monocytes, NK cells and $CD4^+$ T cells. c-e: Mann-Whitney U tests. n=29 d4, n=9 Non-COVID-19 controls. **d-e:** Heat maps of differentially expressed interferon stimulated genes in leukocyte subsets (monocytes and NK cells) of day 4 ambulatory compared to severe hospitalized COVID-19. Monocytes: n=28 upregulated, n=16 downregulated, NK cells: n=25 upregulated, n=15 downregulated. Individual ISG expressions of exemplary ISGs shown. n=29 d4, n=7 hospitalized. **f:** Pearson correlation between the five measured interferon levels and ISG scores of monocytes, NK cells and $CD4^+$ T of day 4 ambulatory COVID-19 patients and hospitalized COVID-19 patients (pooled and day 4 of ambulatory patients alone). P value is shown in center of each field (none <0.05). Pie charts show Pearson's r from maximum 1 (clockwise and blue) to -1 (anticlockwise and red). n=27 day 4 and n=7 hospitalized COVID-19.

**Suppl Fig 8 | Graphical abstract. Upper left:** Schematic of the combined patient cohorts used. n=12 controls, n=46 non-pneumonic/ambulatory SARS-CoV-2 infected patients and n=14 pneumonic/hospitalized COVID-19 patients were included in this study. **Upper right:** Schematic of experimental setup. The study was divided into an exploratory cohort (n=14 patients) using scRNA-Seq and multidimensional flow cytometry of PBMCs and shotgun plasma proteomics. The confirmation cohort (n=58 patients) was used to validate findings from the exploratory cohort, using in-depth RNA-Seq of FACS-sorted PBMCs and multiplex plasma cytokine profiling. **Bottom:** Explanation of findings. After infection with SARS-CoV-2, the virus is either contained in the upper airway tract ("non-pneumonic SARS-CoV-2 infection") or it disseminates into the lung ("pneumonic COVID-19"). Our study showhs that non-pneumonic SARS-CoV-2 infection is characterized by an early strong interferon-stimulated-gene (ISG) signature, as well as an immune regulatory lymphocyte signature and pro-resolving monocytes in the peripheral blood. In contrast, in case of viral dissemination, pneumonic COVID-19 is characterized by lymphocyte cytotoxicity and a proinflammatory marker profile in the peripheral blood.

**Suppl Fig 1**

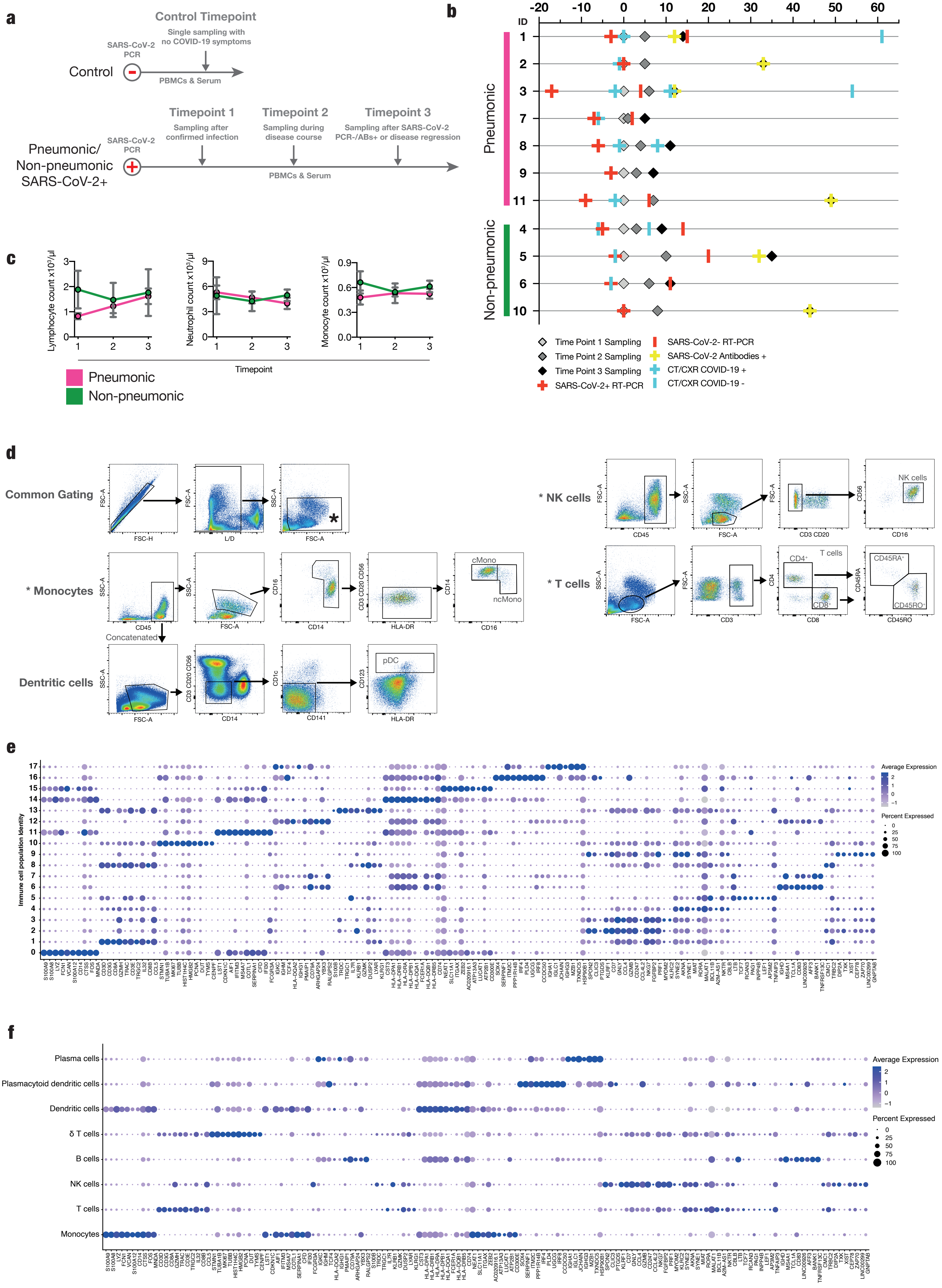
a
Control Timepoint
Single sampling with
no COVID-19 symptoms
SARS-CoV-2
PCR
Control
PBMCs & Serum
Timepoint 1
Sampling after
confirmed infection
Timepoint 2
Sampling during
disease course
Timepoint 3
Sampling after SARS-CoV-2
PCR-/ABs+ or disease regression
Pneumonic/
Non-pneumonic
SARS-CoV-2+
PBMCs & Serum
b
Days post first sampling
ID
Pneumonic
Non-pneumonic
Time Point 1 Sampling
Time Point 2 Sampling
Time Point 3 Sampling
SARS-CoV-2+ RT-PCR
SARS-CoV-2- RT-PCR
SARS-CoV-2 Antibodies +
CT/CXR COVID-19 +
CT/CXR COVID-19 -
c
Lymphocyte count x10³/µl
Neutrophil count x10³/µl
Monocyte count x10³/µl
Timepoint
Pneumonic
Non-pneumonic
d
Common Gating
* Monocytes
Concatenated
Dentritic cells
* NK cells
* T cells
NK cells
cMono
ncMono
pDC
T cells
e
Immune cell population identity
Average Expression
Percent Expressed
f
Plasma cells
Plasmacytoid dendritic cells
Dendritic cells
δ T cells
B cells
NK cells
T cells
Monocytes
Average Expression
Percent Expressed

Suppl Fig 2

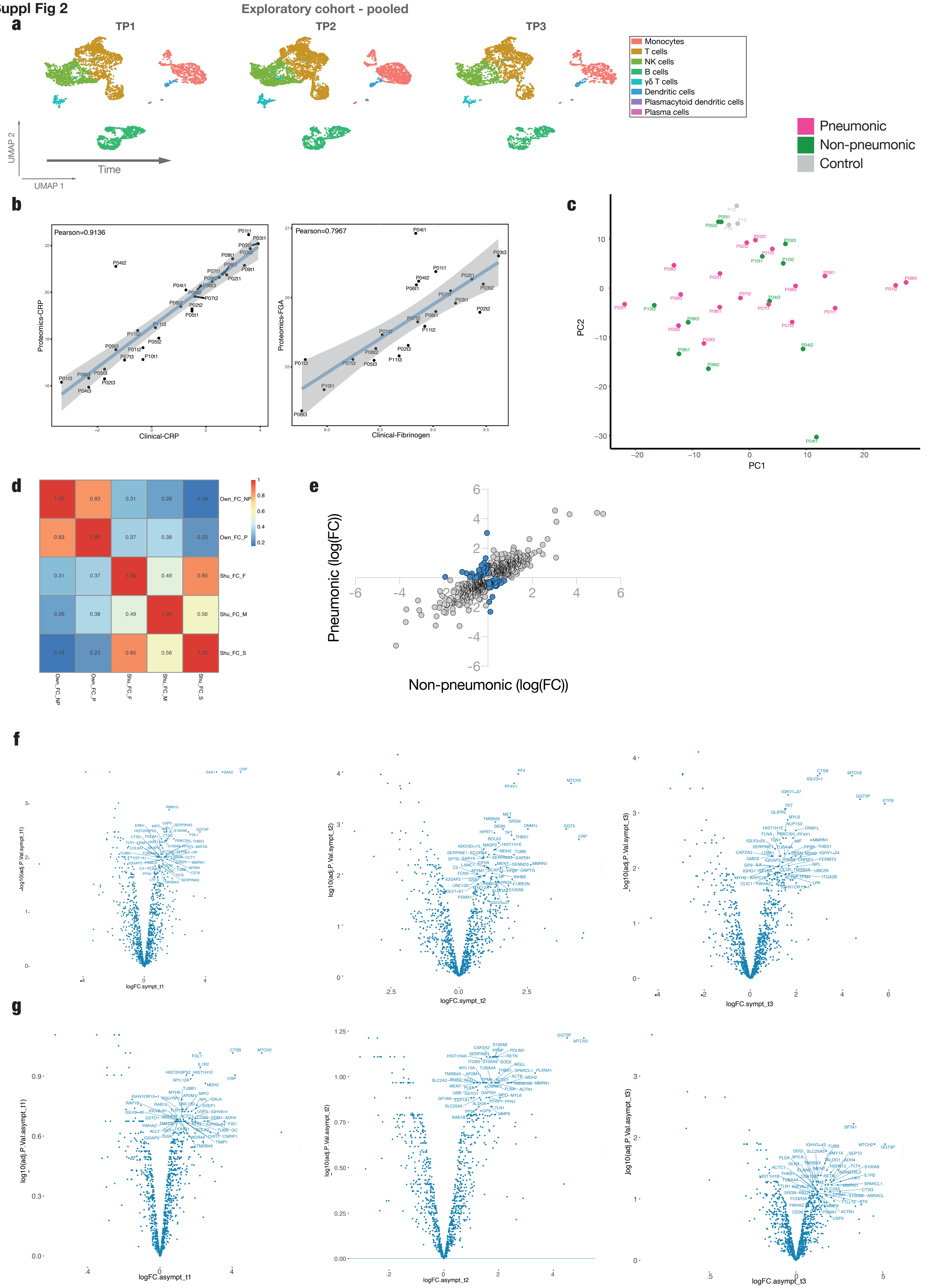
Exploratory cohort - pooled
a
TP1
TP2
TP3
Monocytes
T cells
NK cells
B cells
γδ T cells
Dendritic cells
Plasmacytoid dendritic cells
Plasma cells
UMAP 2
UMAP 1
Time
Pneumonic
Non-pneumonic
Control
b
Pearson=0.9136
Proteomics-CRP
Clinical-CRP
Pearson=0.7967
Proteomics-FGA
Clinical-Fibrinogen
c
PC1
PC2
d
Own_FC_NP
Own_FC_P
Shu_FC_F
Shu_FC_M
Shu_FC_S
e
Pneumonic (log(FC))
Non-pneumonic (log(FC))
f
-log10(adj.P.Val.sympt_t1)
logFC.sympt_t1
log10(adj.P.Val.sympt_t2)
logFC.sympt_t2
log10(adj.P.Val.sympt_t3)
logFC.sympt_t3
g
log10(adj.P.Val.asympt_t1)
logFC.asympt_t1
log10(adj.P.Val.asympt_t2)
logFC.asympt_t2
log10(adj.P.Val.asympt_t3)
logFC.asympt_t3

Suppl Fig 3

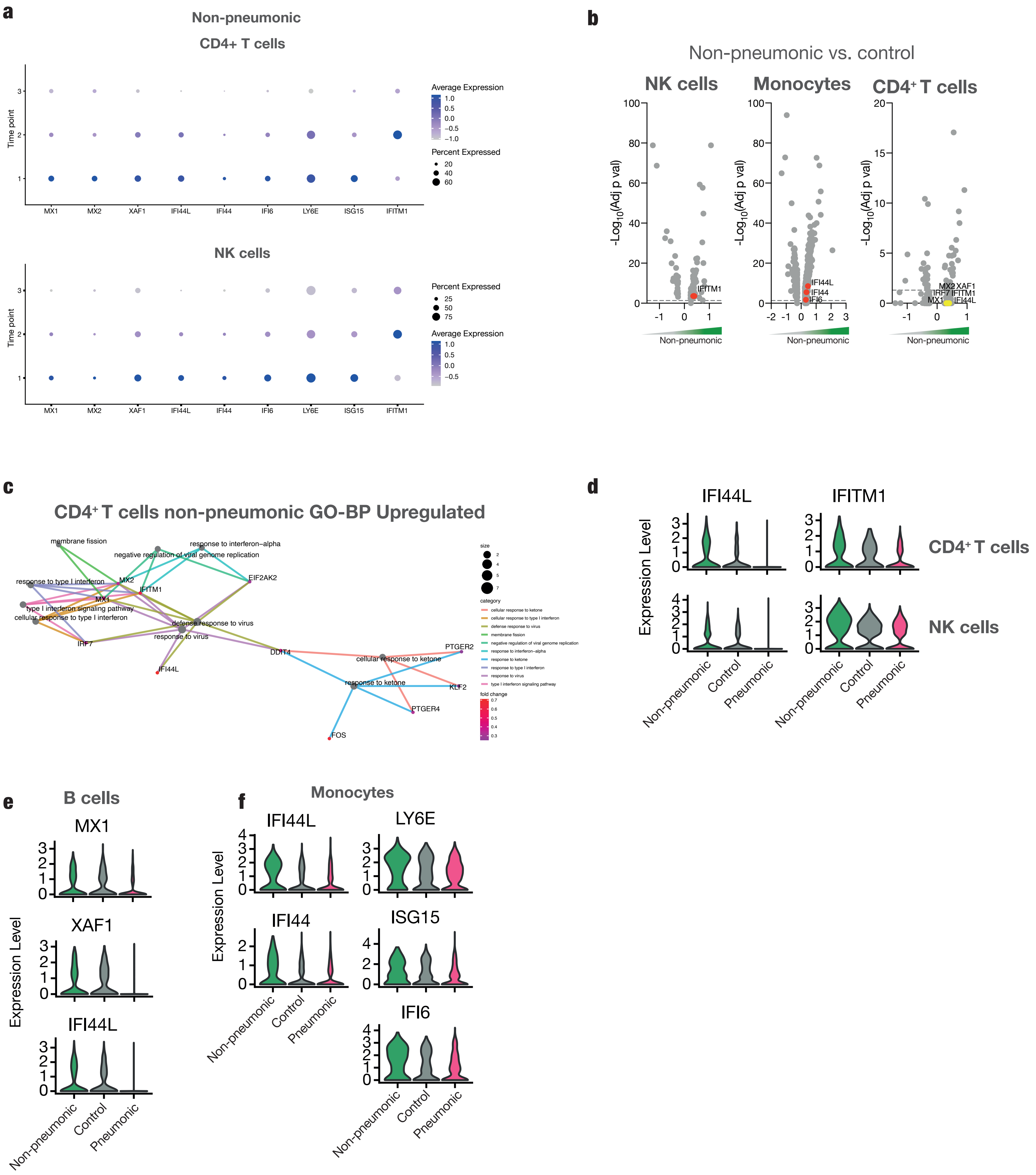
a
Non-pneumonic
CD4+ T cells
Time point
Average Expression
1.0
0.5
0.0
−0.5
−1.0
Percent Expressed
20
40
60
MX1
MX2
XAF1
IFI44L
IFI44
IFI6
LY6E
ISG15
IFITM1
NK cells
Percent Expressed
25
50
75
b
Non-pneumonic vs. control
NK cells
Monocytes
CD4+ T cells
-Log10(Adj p val)
IFITM1
IFI44L
IFI44
IFI6
MX2
XAF1
IRF7
IFITM1
MX1
IFI44L
Non-pneumonic
c
CD4+ T cells non-pneumonic GO-BP Upregulated
membrane fission
response to interferon−alpha
negative regulation of viral genome replication
response to type I interferon
MX2
IFITM1
EIF2AK2
MX1
type I interferon signaling pathway
cellular response to type I interferon
defense response to virus
response to virus
IRF7
IFI44L
DDIT4
PTGER2
cellular response to ketone
response to ketone
KLF2
PTGER4
FOS
size
category
cellular response to ketone
cellular response to type I interferon
defense response to virus
membrane fission
negative regulation of viral genome replication
response to interferon−alpha
response to ketone
response to type I interferon
response to virus
type I interferon signaling pathway
fold change
d
IFI44L
IFITM1
Expression Level
CD4+ T cells
NK cells
Non-pneumonic
Control
Pneumonic
e
B cells
MX1
XAF1
IFI44L
Expression Level
Non-pneumonic
Control
Pneumonic
f
Monocytes
IFI44L
LY6E
IFI44
ISG15
IFI6
Expression Level
Non-pneumonic
Control
Pneumonic

Suppl Fig 4

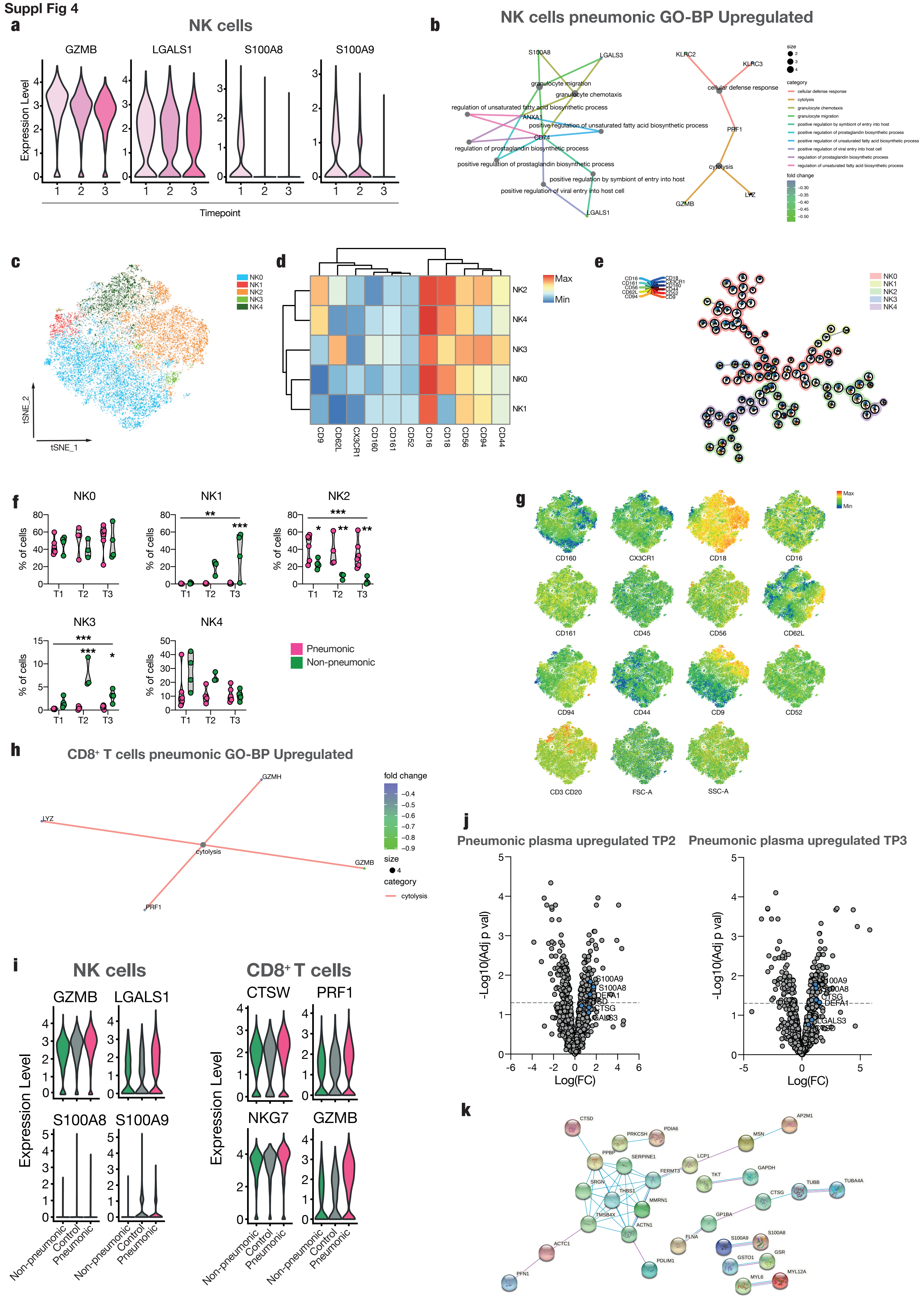
a
NK cells
GZMB
LGALS1
S100A8
S100A9
Expression Level
Timepoint
b
NK cells pneumonic GO-BP Upregulated
c
NK0
NK1
NK2
NK3
NK4
tSNE_2
tSNE_1
d
Max
Min
e
f
NK0
NK1
NK2
NK3
NK4
% of cells
T1
T2
T3
Pneumonic
Non-pneumonic
g
CD160
CX3CR1
CD18
CD16
CD161
CD45
CD56
CD62L
CD94
CD44
CD9
CD52
CD3 CD20
FSC-A
SSC-A
h
CD8+ T cells pneumonic GO-BP Upregulated
GZMH
LYZ
cytolysis
GZMB
PRF1
fold change
size
category
i
NK cells
CD8+ T cells
CTSW
PRF1
NKG7
Non-pneumonic
Control
Pneumonic
j
Pneumonic plasma upregulated TP2
Pneumonic plasma upregulated TP3
-Log10(Adj p val)
Log(FC)
k

Suppl Fig 5

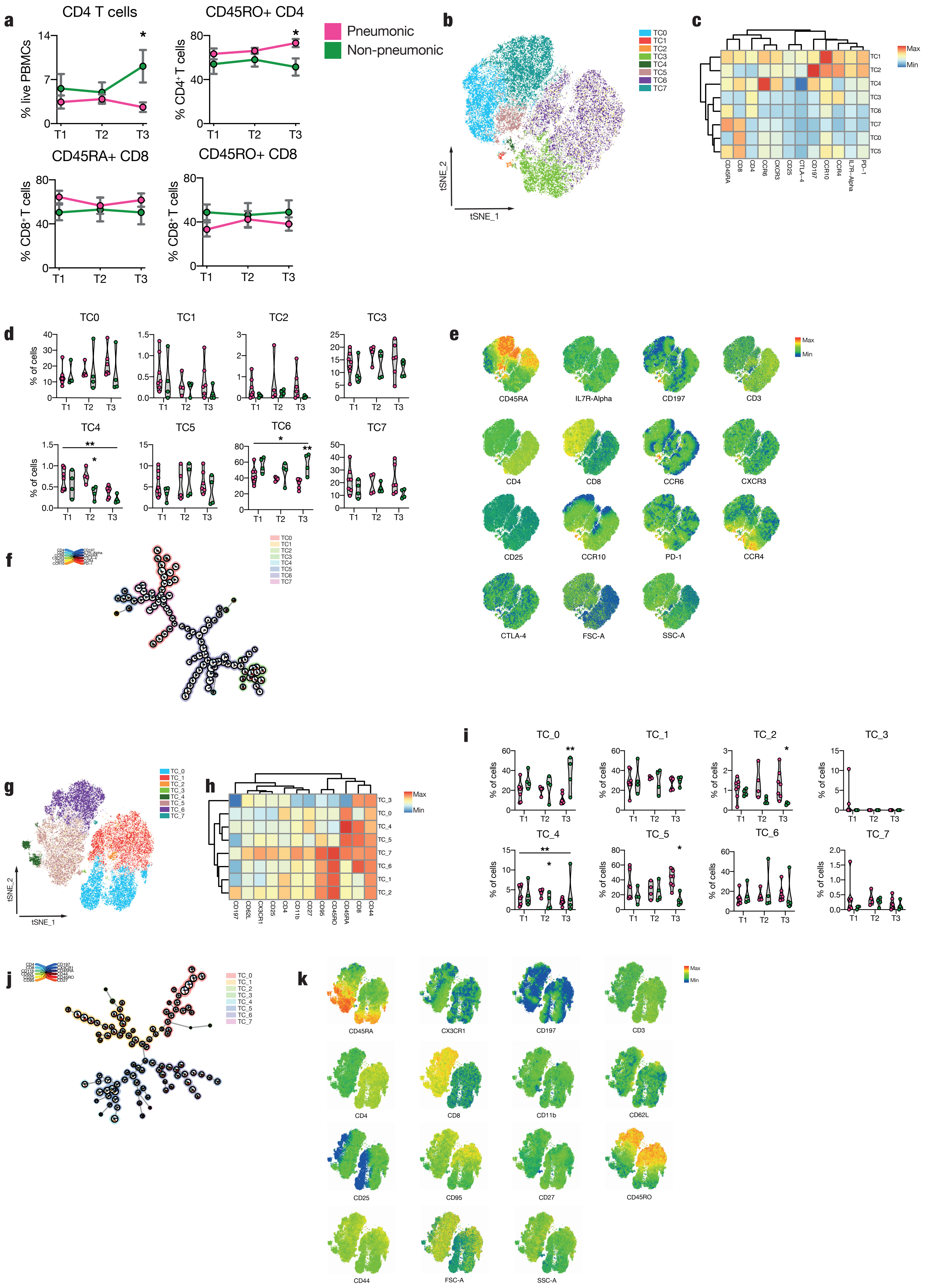
a
CD4 T cells
CD45RO+ CD4
CD45RA+ CD8
CD45RO+ CD8
% live PBMCs
% CD4⁺ T cells
% CD8⁺ T cells
T1
T2
T3
Pneumonic
Non-pneumonic
b
TC0
TC1
TC2
TC3
TC4
TC5
TC6
TC7
tSNE_1
tSNE_2
c
Max
Min
CD45RA
CD8
CD4
CCR6
CXCR3
CD25
CTLA-4
CD197
CCR10
CCR4
IL7R-Alpha
PD-1
d
% of cells
e
CD45RA
IL7R-Alpha
CD197
CD3
CD4
CD8
CCR6
CXCR3
CD25
CCR10
PD-1
CCR4
CTLA-4
FSC-A
SSC-A
f
g
TC_0
TC_1
TC_2
TC_3
TC_4
TC_5
TC_6
TC_7
h
CD197
CD62L
CX3CR1
CD25
CD4
CD11b
CD27
CD95
CD45RO
CD45RA
CD8
CD44
i
j
k
CD45RA
CX3CR1
CD197
CD3
CD4
CD8
CD11b
CD62L
CD25
CD95
CD27
CD45RO
CD44
FSC-A
SSC-A

Suppl Fig 6

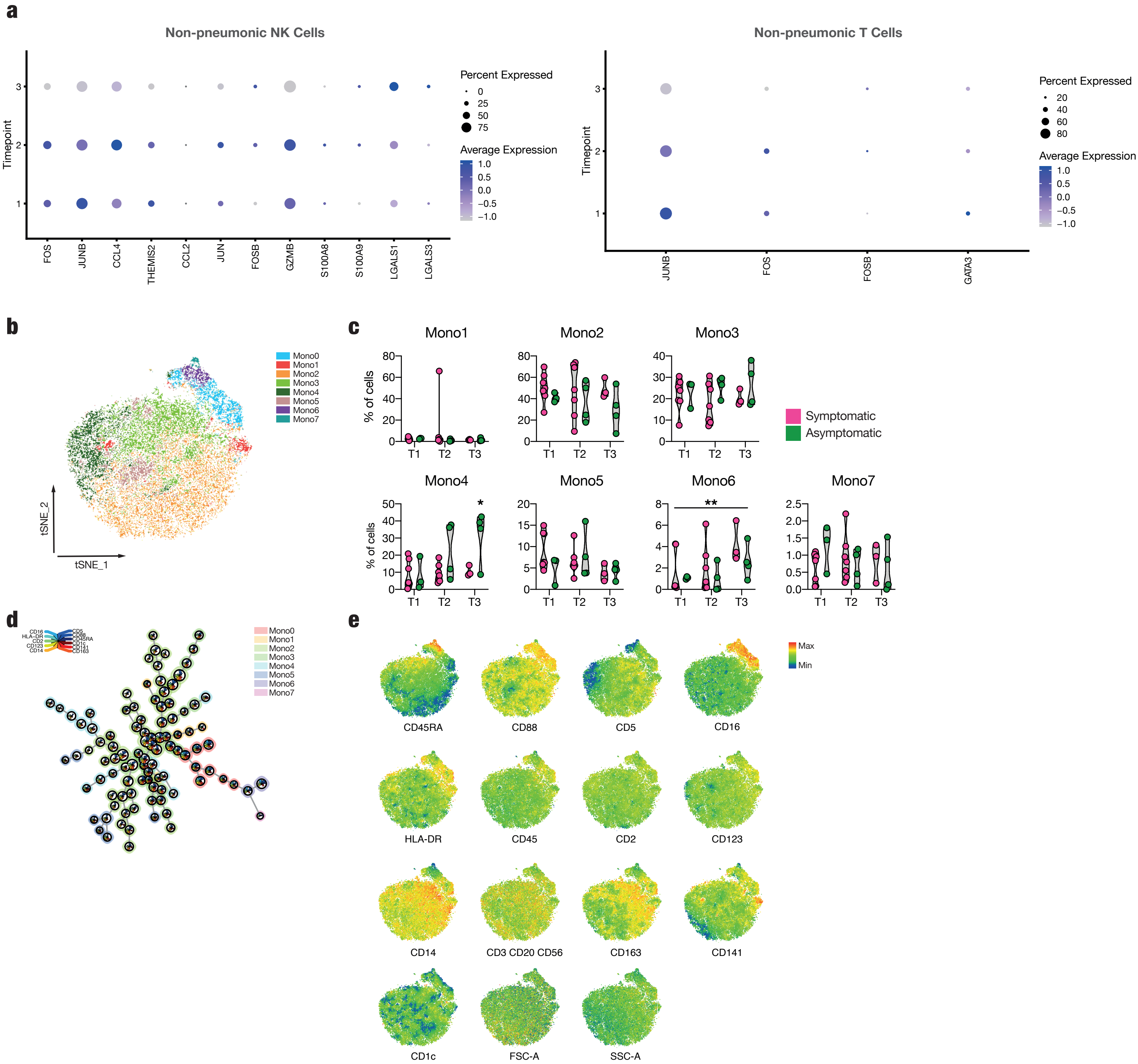
a
Non-pneumonic NK Cells
Timepoint
3
2
1
FOS
JUNB
CCL4
THEMIS2
CCL2
JUN
FOSB
GZMB
S100A8
S100A9
LGALS1
LGALS3
Percent Expressed
0
25
50
75
Average Expression
1.0
0.5
0.0
−0.5
−1.0
Non-pneumonic T Cells
Timepoint
3
2
1
JUNB
FOS
FOSB
GATA3
Percent Expressed
20
40
60
80
Average Expression
1.0
0.5
0.0
−0.5
−1.0
b
Mono0
Mono1
Mono2
Mono3
Mono4
Mono5
Mono6
Mono7
tSNE_2
tSNE_1
c
Mono1
Mono2
Mono3
Mono4
Mono5
Mono6
Mono7
% of cells
T1
T2
T3
*
**
Symptomatic
Asymptomatic
d
CD16
HLA-DR
CD2
CD123
CD14
CD5
CD88
CD45RA
CD1c
CD141
CD163
Mono0
Mono1
Mono2
Mono3
Mono4
Mono5
Mono6
Mono7
e
Max
Min
CD45RA
CD88
CD5
CD16
HLA-DR
CD45
CD2
CD123
CD14
CD3 CD20 CD56
CD163
CD141
CD1c
FSC-A
SSC-A

Suppl Fig 7

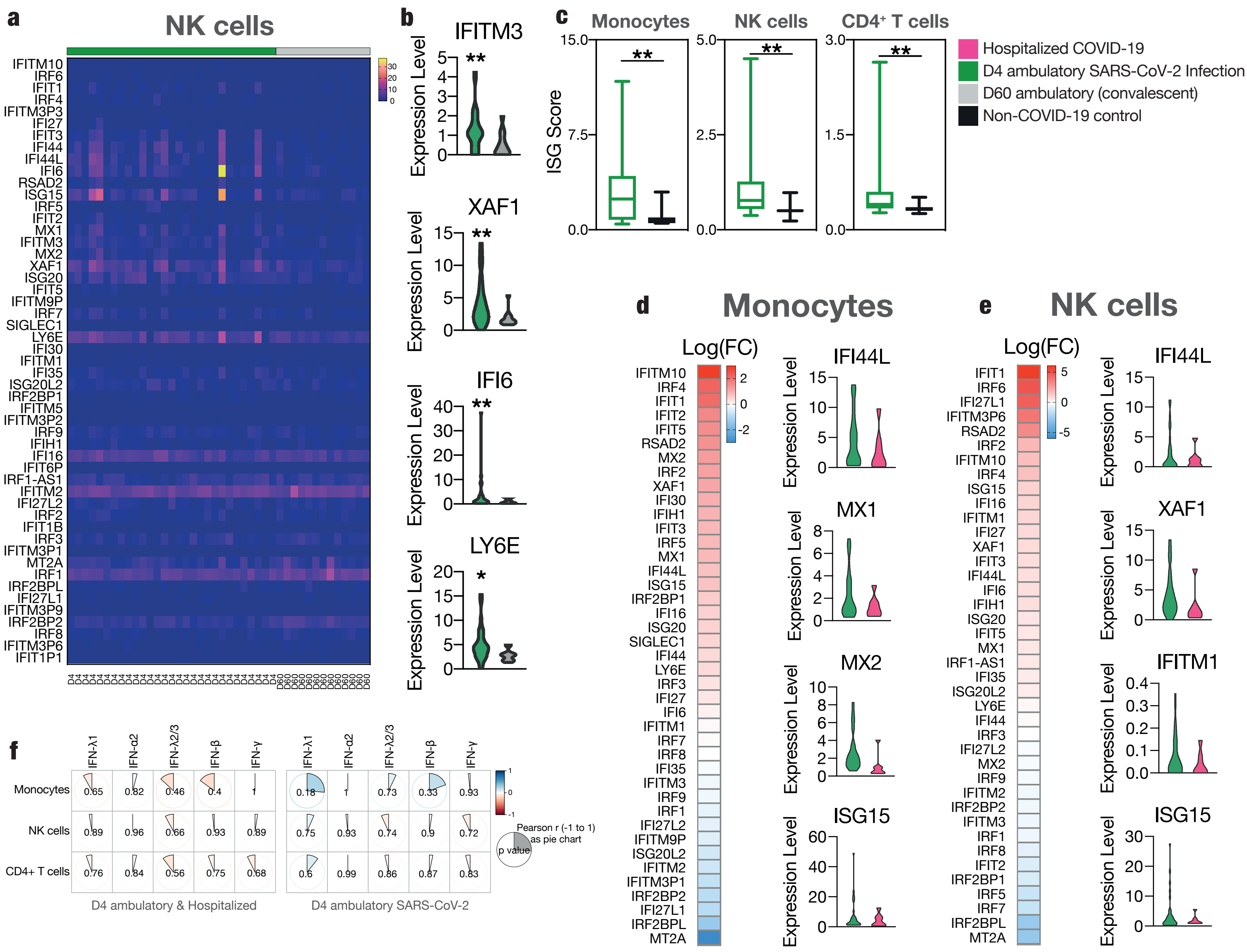

a
NK cells
b
IFITM3
XAF1
IFI6
LY6E
Expression Level
c
Monocytes
NK cells
CD4+ T cells
ISG Score
Hospitalized COVID-19
D4 ambulatory SARS-CoV-2 Infection
D60 ambulatory (convalescent)
Non-COVID-19 control
d
Monocytes
Log(FC)
IFI44L
MX1
MX2
ISG15
e
NK cells
Log(FC)
IFI44L
XAF1
IFITM1
ISG15
f
IFN-λ1
IFN-α2
IFN-λ2/3
IFN-β
IFN-γ
Monocytes
NK cells
CD4+ T cells
Pearson r (-1 to 1) as pie chart
p value
D4 ambulatory & Hospitalized
D4 ambulatory SARS-CoV-2

Suppl Fig 8

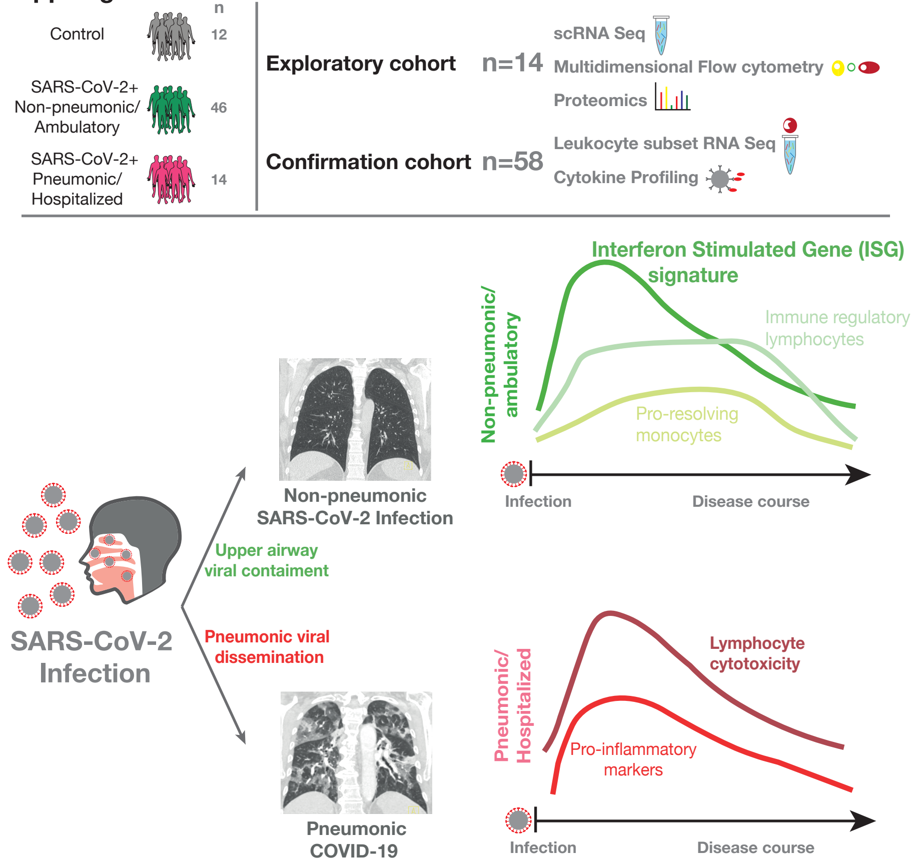

n
Control
12
SARS-CoV-2+
Non-pneumonic/
Ambulatory
46
SARS-CoV-2+
Pneumonic/
Hospitalized
14
Exploratory cohort
n=14
scRNA Seq
Multidimensional Flow cytometry
Proteomics
Confirmation cohort
n=58
Leukocyte subset RNA Seq
Cytokine Profiling
Interferon Stimulated Gene (ISG)
signature
Immune regulatory
lymphocytes
Pro-resolving
monocytes
Non-pneumonic/
ambulatory
Infection
Disease course
Non-pneumonic
SARS-CoV-2 Infection
Upper airway
viral containment
SARS-CoV-2
Infection
Pneumonic viral
dissemination
Pneumonic
COVID-19
Pneumonic/
Hospitalized
Lymphocyte
cytotoxicity
Pro-inflammatory
markers
Infection
Disease course
